# Structure and function of a novel osmoregulated periplasmic fiber-forming high-molecular-weight carbohydrate of *Myxococcus xanthus*

**DOI:** 10.1101/2020.07.26.208595

**Authors:** Sapeckshita Agrawal, Christian Heiss, David M. Zuckerman, Jeffery M. T. So, Koen Semeijn, Radnaa Naran, Parastoo Azadi, Egbert Hoiczyk

**Affiliations:** Department of Molecular Microbiology and Immunology, Johns Hopkins Bloomberg School of Public Health, 605 North Wolfe Street, Baltimore, MD 21205, USA; Complex Carbohydrate Research Center, The University of Georgia, 315 Riverbend Rd., Athens, GA 30602; Department of Molecular Biology and Biotechnology, The Krebs Institute, University of Sheffield, Firth Court, Western Bank, Sheffield, S10 2TN, UK; Iona College, 715 North Avenue, New Rochelle, NY 10801

**Keywords:** High-molecular-weight carbohydrate, Sugar analysis, Periplasmic glucans, Periplasmic space, Osmoregulation, Osmolyte, Turgor, Spore formation, *Myxococcus xanthus*

## Abstract

Osmoregulation is of central importance for living cells. In Gram-negative bacteria, strategies for osmoregulation and turgor maintenance in hypotonic environments include the synthesis, accumulation, and modification of periplasmic oligosaccharides. These osmoregulated periplasmic glucans (OPGs, formerly known as membrane-derived oligosaccharides or MDOs) promote water uptake and retention, keeping the cells in an optimal state of hydration. While our understanding of OPG-dependent osmoregulation in a number of model organisms like *Escherichia coli* is quite detailed, less is known about these processes in bacteria that live in environments characterized by strongly fluctuating osmolarity, such as soil. Here we describe that the soil bacterium *Myxococcus xanthus* lacks a canonical low-molecular-weight OPG, but instead possesses a novel high-molecular-weight, fiber-forming polysaccharide. Chemical analysis reveals that this polysaccharide is several thousand kilodaltons in size, composed of a highly branched decasaccharide repeat unit containing mannose, glucose, N-acetylglucosamine, and rhamnose. Physiological experiments indicate that the polysaccharide is osmoregulated thereby functionally replacing the canonical OPG. Moreover, experiments indicate that this high-molecular-weight periplasmic polysaccharide forms a fibrillar meshwork that stabilizes the cell envelope during glycerol spore formation, a process during which the entire peptidoglycan of the cell is degraded and the rod-shaped vegetative cells convert into spherical spores.

**Significance:** Osmoprotection is a necessity for every living cell, particularly in an environment with fluctuating osmolarity. In Gram-negative bacteria, low-molecular-weight osmoregulated periplasmic glucans (OPGs) are an important component of the osmotic stress response in hypotonic environments. Here, we describe that the soil bacterium *Myxococcus xanthus* does not possess such an OPG but instead accumulates a novel high-molecular-weight fiber-forming polysaccharide in the periplasm in response to hypotonic conditions. This polymer is important for osmoprotection of the cells and plays a key role in the stabilization of the cell envelope during the conversion of rod-shaped vegetative cells into spherical spores. These results indicate that bacteria may use non-OPG carbohydrates for osmoprotection and cell wall stabilization during processes like cellular differentiation.

## Introduction

Maintaining a stable cell turgor is essential for bacteria to sustain rigidity, control transport processes and solute concentrations, and allow for normal growth (Wood, 2011). Events that disturb this balance induce physiological responses that allow cells to re-establish their turgor and prevent or minimize potentially harmful consequences. In all proteobacteria examined, low environmental osmolarity triggers the synthesis and accumulation of osmoregulated periplasmic glucans (OPGs), polyglucose oligosaccharides that are an important component of the periplasm (Schulman and Kennedy, 1979; Ehrmann, 2006). OPGs were first discovered in the culture filtrates of *Agrobacterium tumefaciens* and initially mistaken as low-molecular-weight (MW) exopolysaccharides (EPSs) (McIntire et al., 1942). With the subsequent discovery of the “membrane-derived oligosaccharides” (MDOs) in *Escherichia coli,* a second group of OPGs was identified (Van Golde et al., 1973), and it was eventually recognized that both the cyclic glucans of *Agrobacterium* and the linear glucans of *Escherichia* are periplasmic osmoprotective carbohydrates (Kennedy, 1982; Miller et al., 1986). Since then, research has shown that all characterized OPGs share a number of core features (reviewed in Bohin, 2000 and Lee at al., 2009 and references therein). They all are short, 5-40 glucose unit-long oligosaccharides that are mostly linked via β-glycosidic bonds and synthesized in the periplasm in response to low environmental osmolarity. Based on their size and the linkage of their polyglucan backbone, OPGs have been classified into four different families. Family I, which is found in *Escherichia*, *Erwinia,* and *Pseudomonas* species is heterogeneous in length ranging from 5-28 glycosyl residues that form a β-1,2-linked linear backbone carrying β-1,6-linked branches of glycosyl units. Family II, which has been described in *Agrobacterium*, *Sinorhizobium*, *Mesorhizobium*, and *Brucella* species, form cyclic structures by β-1,2-linkages of glycans that are 17-40 glycosyl residues long. Like family II, family III OPGs are cyclic, but in contrast they are shorter, only 10-13 glycosyl units long, and possess β-1,3 and β-1,6 glycosidic bonds. OPGs of this family have so far been found in *Bradyrhizobium*, *Azorhizobium*, and *Azospirillum* species. Finally, family IV OPGs, described in *Xanthomonas*, *Ralstonia*, and *Rhodobacter* species, are cyclic, 13-18 glycosyl unit-long oligosaccharides that contain a β-1,2-linkage backbone with one additional single α-1,6-linkage. OPGs of all four families can also contain additional non-sugar substituents. Two types of such substituents have been identified: molecules which derive from the turnover of phospholipids such as phosphoethanolamine, phosphoglycerol, and phosphocholine, and substituents that are products of the cellular metabolism like acetyl, succinyl, and methylmalonyl residues (Bohin, 2000; Lee et al., 2009). Both the nature and the degree of substitution can vary within a species, reflecting the growth rate of the bacteria as well as environmental conditions. For example, the family II type cyclic OPG from *Sinorhizobium meliloti* can be converted from a neutral molecule to a highly charged anionic oligomer, a conversion that is more rapid and complete in exponentially growing cells than in stationary cultures (Geiger et al., 1991). As some OPGs are never substituted, the overall role of substitutions for the physiological function of OPGs is not well understood. Although there is a great deal of diversity in the composition of OPGs, it is generally accepted that all OPGs play a major role in osmoprotection by keeping the periplasms of cells hydrated (Kennedy, 1982; Miller et al., 1986). Beyond this osmoprotective function, OPGs have been proposed to play roles in a few well studied bacteria in virulence, host colonization, cell signaling, protein folding, and envelope structure (reviewed in Bontemps-Gallo at al. 2017). While OPGs have been studied in many α-, β-, and γ-proteobacteria, in ε-proteobacteria so far only functional analogs, called OPG-like oligosaccharides, have been identified, and no information exists regarding OPGs in ξ- and δ-proteobacteria (Bontemps-Gallo and Lacroix, 2015).

*Myxococcus xanthus* is a common soil-dwelling myxobacterium of the δ-proteobacteria class that is highly social and forms swarms that cooperatively prey on other bacteria (Berleman and Kirby, 2009). Possessing one of the most complex prokaryotic genomes (Goldman et al., 2006), these bacteria perform surface motility (Chang at al. 2016; Faure et al., 2016) and, upon starvation, initiate a complex developmental program during which vegetative cells aggregate and a subset of cells undergo a rod-to-sphere transition to form spores within a fruiting body (Zusman et al., 2007). Like all soil organisms, *M. xanthus* has developed adaptations that help the cells cope with their frequently fluctuating environment. One behavioral strategy is developmental differentiation that allows the bacteria to match cell density with available food resources (Berleman et al., 2008). Another strategy is phase switching, a genetic program that generates phenotypic variation through the expression of different sets of genes in subpopulations of bacteria (Furusawa et al., 2011; Higgs et al., 2014). This variation provides a selective advantage because members of a subpopulation will be better adapted to survive under certain environmental conditions (Higgs et al., 2014; Dziewanoswka et al., 2014). While both strategies, differentiation and phase switching, work on the population level, much less is known about how individual cells physiologically cope with changes in the environment such as water availability. Here, we describe that *M. xanthus* lacks a canonical OPG under standard laboratory growth conditions, but instead synthesizes a periplasmic high-MW carbohydrate that appears to protect the cells during developmental cell-to-spore transition. This novel carbohydrate is many hundred times larger in size than any previously described OPGs, and fibers are so large that they are visible by electron microscopy. Moreover, the polymer also differs chemically from known OPGs, containing several sugars other than glucose. Despite these differences, however, it performs a physiological function comparable to OPGs, as its synthesis is osmoregulated. Taken together, these data indicate that bacteria living in environments characterized by wide ranging osmotic fluxes may use unusual polymeric high-MW OPGs that have so far been overlooked as they are not liberated upon classic OPG isolation procedures.

## Results

### *Myxococcus xanthus* synthesizes a high-MW carbohydrate

During attempts to isolate pili, we noted that when *M. xanthus* cells were subjected to high shear forces, but not broken, the supernatant of the cell suspension became highly viscous (Fig. 1A). Since efforts to lower this viscosity through digestion with DNase failed, we concluded that a non-nucleic acid polymer had been released; an interpretation that was supported using the carbohydrate-specific anthrone test (Seifter et al., 1950). Taking advantage of its high MW (Fig. 1B), we developed a purification scheme for this carbohydrate (for details see Materials and Methods). To exclude the possibility that this polymer was part of the exopolysaccharide (EPS) matrix of *M. xanthus*, we repeated the isolations using liquid-grown cells of the EPS-deficient ΔdifE strain (Black and Yang, 2004; Lu et al., 2005). Electron microscopy confirmed that the isolated carbohydrate in both cases was a high-MW polymer formed by ca. 4 nm wide filaments that, like cellulose fibers (Hiagler, 1985; Jakob et al., 1995), aggregated into thicker bundles up to 40 nm wide (Fig. 1C). Initial attempts to further purify the polymer using a gel filtration column (molecular cutoff 500 kDa) failed because the native molecule eluted in the void volume, consistent with a molecule of very large size. Upon partial hydrolysis, we could reproducibly isolate a ca. 400 kDa large fragment suitable for chemical analyses (Fig. 1D; and below).

**Fig. 1.**
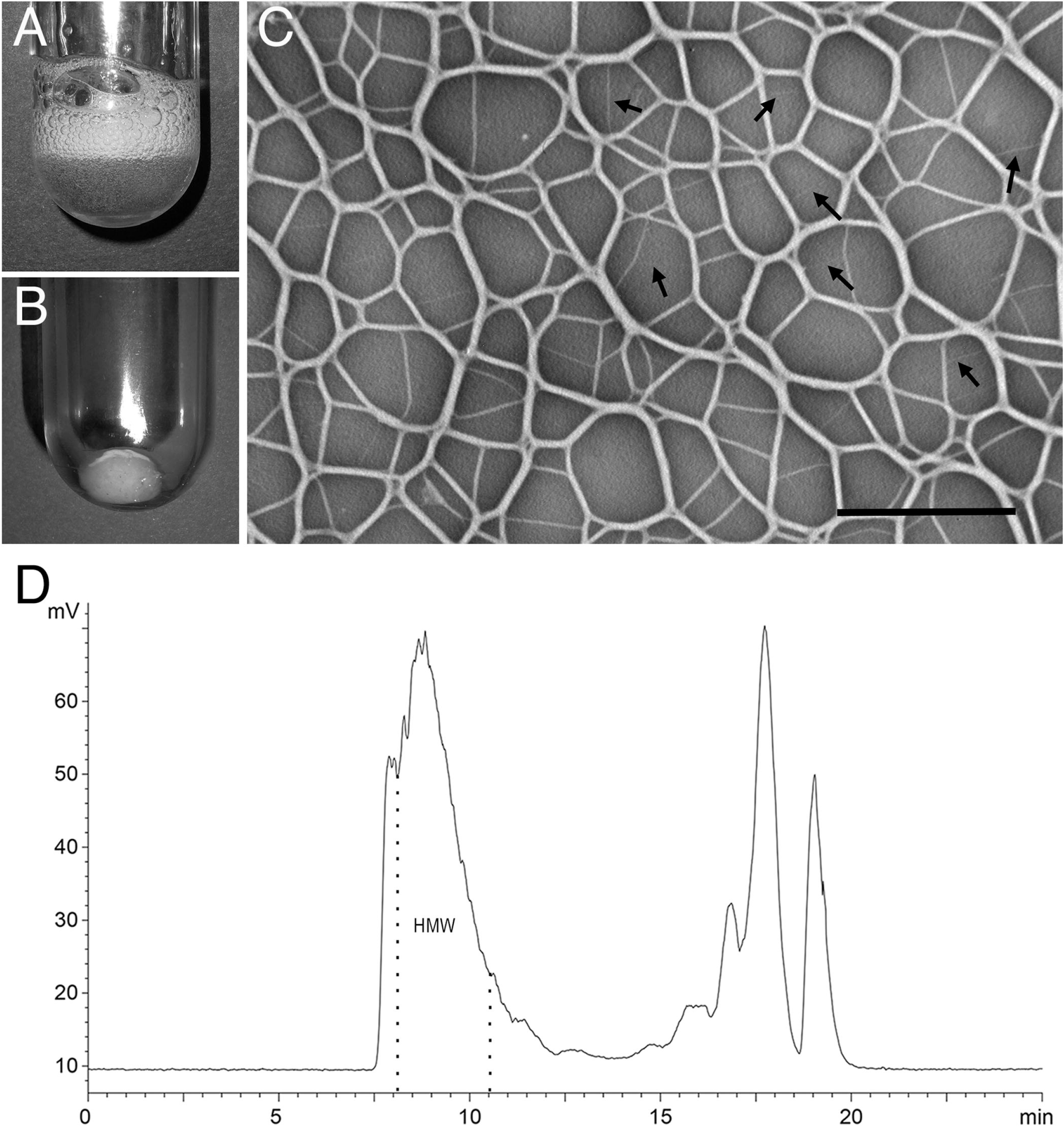
*Myxococcus xanthus* cells produce a high-MW polymer that can be extracted by subjecting the cells to high shear forces. (*A*) The released purified polymer increases the viscosity of water visible through the trapping of bubbles and (*B*) can be pelleted using ammonium sulfate. (*C*) In negatively stained preparations, the isolated polymer is composed of thin fibrils ca. 4 nm wide that often bundle into larger fibers forming a fibrillar meshwork. Bar, 0.5 μm. (*D*) After partial hydrolysis the molecule fragments and the fraction containing the ca. 400 kDa large fragment (high-molecular-weight, HMW) was isolated and used for chemical analyses.

### The isolated polymer is located in the periplasm of the cell, which lacks a canonical OPG under standard culture conditions

Following the observation that we could recover this carbohydrate from the EPS-deficient ΔdifE strain, strongly suggesting that the material was not part of the EPS matrix, we next attempted to more specifically determine its cellular localization. Polyclonal antibody raised against the purified polymer preferentially labeled the cell envelopes of ΔdifE cells, indicating that the polymer is located either on the surface of the outer membrane or in the periplasm (Fig. 2A). Since our initial observation indicated that the polymer is only released when the cells are subjected to high shear, we next tested various treatments for their ability to release the material (Fig. 2B). By far the most effective treatment was osmotic shock, a procedure that specifically releases the periplasmic content of Gram-negative bacteria (Nossal and Heppel, 1966). Less effective was agitation in a blender, while shaking or washing of the cells with salt solutions, methods usually used for EPS isolation, released only negligible amounts of polymer. Together these results indicated that the outer membrane had to be physically ruptured to release the material, an observation that supported the idea that the polymer is located in the periplasm. To further corroborate this conclusion, we next established that the polymer was released under the same conditions as the well-established periplasmic marker, alkaline phosphatase (Nossal and Heppel, 1966; Fig. 2C, dark grey bars), and that during these extractions the cytoplasmic membrane was not breached, and only negligible amounts of cytoplasmic content released, using the cytoplasmic protein EncA as a marker (McHugh et al., 2014; Fig. 2D). Notable, the measured alkaline phosphatase data match closely data reported in an earlier publication (Rodriguez-Soto and Kaiser, 1997; Fig. 2C, light grey bars). In summary, these results demonstrated that the polymer is periplasmic and only released using methods and conditions that liberate known periplasmic markers.

**Fig. 2.**
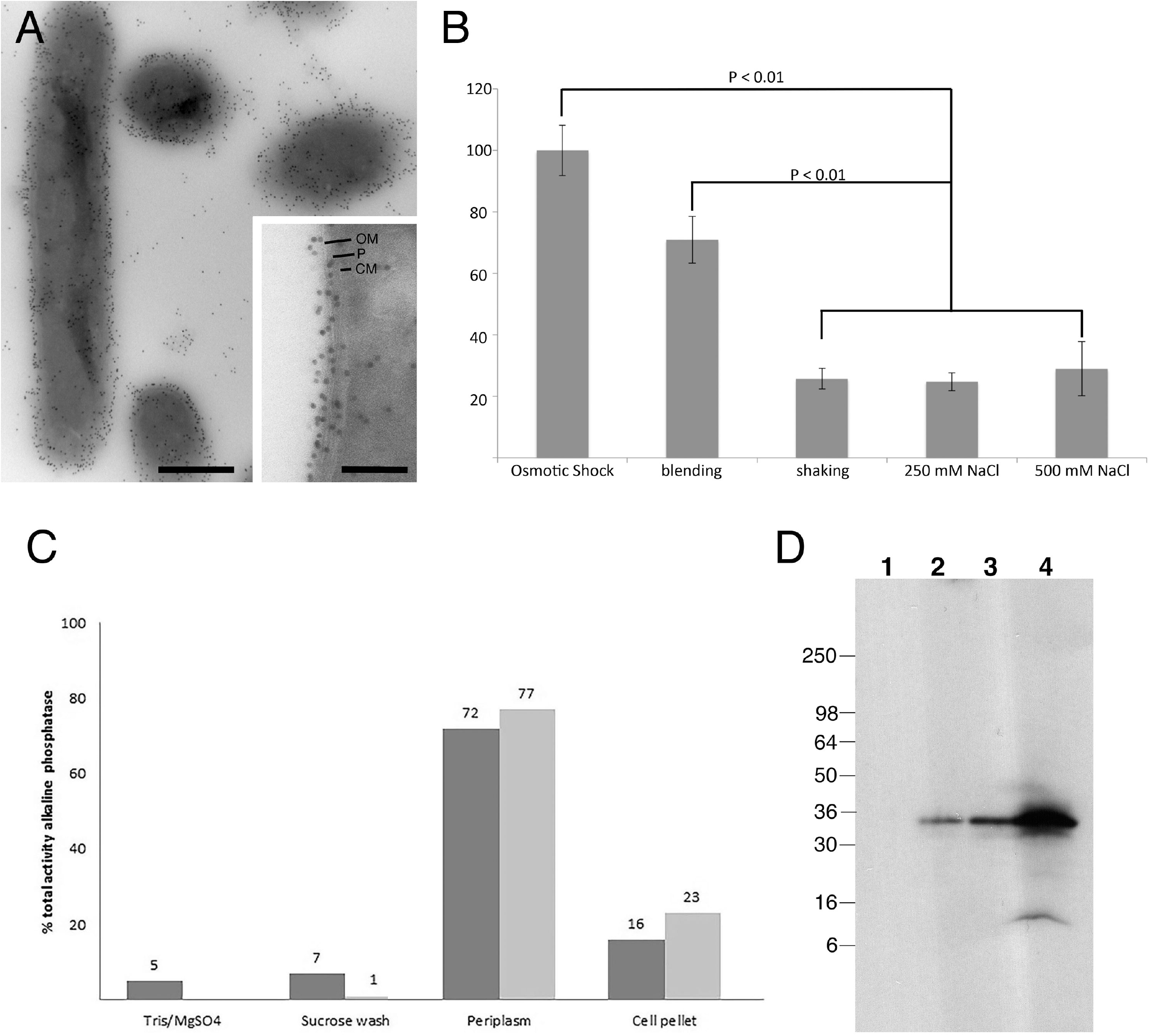
A high-MW polymer localizes to the periplasm of *Myxococcus xanthus.* (*A*) Cryo-immuno electron microscopy reveals that an anti-polymer antibody labels the cell envelope of *ΔdifE* cells in thin sections. (*B*) Detection of the release of the polymer under various treatment regiments. Only treatments that rupture the outer membrane, such as sucrose shock and high-speed blending, release the polymer, whereas procedures aimed at extracting extracellular polysaccharides, such as salt washes, are ineffective. The variance was determined by one-way ANOVA and p-values were calculated with the Tukey test. (*C*) Alkaline phosphatase, a known periplasmic enzyme is released under the same conditions as the polymer. The black bars show the experimental data, while the grey bars are from Rodriguez-Soto and Kaiser, 1997 (no Tris/MgSO_4_ wash was performed by these authors). (*D*) Release of EncA during various cell treatments. The cytoplasmic protein EncA was used to monitor the release of cytoplasmic content during various treatments of cells used to extract the periplasmic polymer. Conditions that breach the outer membrane and release the polymer release only small amounts of EncA, indicating that the cytoplasmic membrane is not substantially breached under these conditions. Lane 1, Tris/MgSO_4_ wash; lane 2, sucrose wash; lane 3, periplasm; lane 4, cell pellet.

Since OPGs have commonly been found in the periplasm of other Gram-negative bacteria (Bontemps-Gallo et al., 2017), we next tested whether *M. xanthus* also contained a canonical low-molecular-weight OPG. *M. xanthus* wt and ΔdifE cells were grown in both liquid culture and on agar, collected, and treated with 5% trichloroacetic acid, a standard method for isolating OPGs. We failed to detect the release of any OPGs, indicating that under standard growth conditions, the cells appear to lack this periplasmic component.

### Chemical analysis shows that the *M. xanthus* polymer is highly branched and formed by a complex decasaccharide repeat unit

Using glycosyl composition analysis of the dialyzed *M. xanthus* periplasmic high-MW carbohydrate (MXP), we identified the monosaccharides constituting the polymer as mannose, glucose, N-acetylglucosamine, and rhamnose. We performed a linkage analysis of MXP using the experimental sequence of permethylation, hydrolysis, reduction, and acetylation (Pettolino et al., 2012), and detected the resulting partially methylated alditol acetates (PMAAs) by gas chromatography with mass spectrometric detection (GC-MS). We found PMAAs corresponding to the following major linkages (Table S1): terminal, 4-linked, and 2,3-linked rhamnose (Rha), terminal, 3-linked, and 2,3,4-linked glucose (Glc), 3-linked and 2,3-linked mannose (Man), and terminal and 3-linked N-acetylglucosamine (GlcNAc). Aqueous solutions of the native MXP were too viscous to allow acquisition of useful NMR spectra. For this reason, we treated the MXP with dilute acid to obtain a more tractable material, which was isolated by semi-preparative size exclusion high performance liquid chromatography (SEC-HPLC). The resulting size-reduced polysaccharide (MXP-A) had an average molecular weight of 400 kDa and was much more amenable to solution-state NMR analysis. We acquired a series of 1- and 2-dimensional NMR spectra of MXP-A (Fig. S3) and found that the spectra were remarkably complex, showing more than 25 anomeric signals. To simplify the structure further, we performed a Smith degradation on MXP-A (Perlin, 2006; MacLean and Perry, 2010). We separated the resulting products by SEC-HPLC and detected both polymeric material and released monosaccharides, collecting the tail end of the polymeric peak for further characterization. Linkage analysis of the Smith degraded polymer (MXP-AS) (Table S1) showed that the 2,3,4-Glc and 3-GlcNAc linkages were still present, while the 2,3-Man, the 2,3-Rha, and the 4-Rha linkages had disappeared. Although terminal residues do not survive the Smith degradation, among the terminal sugars in the original material, only terminal GlcNAc disappeared in the linkage analysis of MXP-AS. We interpret this to mean that new terminal Glc and Rha residues were created from the previous 2,3-Rha and 3-Glc. The proton NMR spectrum showed 5 major anomeric signals, 2 prominent signals near 4.2 ppm, consistent with the H-2 proton of Man or Rha, a complex carbohydrate bulk region, an N-acetyl signal near 2 ppm, and a Rha H-6 signal near 1.3 ppm. Following the connectivities in the proton-proton Correlation Spectroscopy (COSY) and Total Correlation Spectroscopy (TOCSY) spectra and reading carbon chemical shifts from the proton-carbon Heteronuclear Single Quantum Correlation (HSQC) spectrum (Fig. S4) (Duus et al., 2000) allowed us to assign nearly all of the chemical shifts of the 5 major monosaccharide residues present in the MXP-AS sample (Table S2) and to identify sugar type and anomeric configuration. Thus, the 5 residues were identified as 2,3,4-α-Glc (Residue **A**), 3-α-Man (Residue **B**), β-3-GlcNAc (Residue **C**), α-Rha (Residue **D**), and α-Glc (Residue **E**), all in pyranose form. Proton-proton Nuclear Overhauser Effect Spectroscopy (NOESY) and proton-carbon Heteronuclear Multiple Bond Correlation (HMBC) spectra (Fig. S4) were used to determine the connectivities between these residues and to define the sequence of MXP-AS. NOESY cross peaks were observed between H-1 of **A** and H-3 of **B**, H-1 of **B** and H-3 of **C**, H-1 of **D** and H-4 of **A**, and H-1 of **E** and H-2 of **A**. Only H-1 of **C** did not show any inter-residue NOE cross peak. However, the HMBC spectrum revealed the connectivity of Residue **C** by displaying a cross peak between its H-1 and C-3 of Residue **A**. The HMBC spectrum also confirmed the connectivities of the other residues. Taken together, these data lead to the conclusion that MXP-AS has a pentasaccharide repeating unit consisting of a [3-β-GlcNAc-1-3-α-Glc-1-3-α-Man-1-] trisaccharide backbone, decorated with α-Glc on O-2 and α-Rha on O-4 of the backbone Glc residue (Fig 3A). All residues were in pyranose form (omitted in the structure drawing for simplicity’s sake). Having thus determined the core structure of MXP, we set out to establish the location of the residues that were removed during the Smith degradation. Comparing the linkage data of MXP and MXP-AS, we were able to determine the positions that were glycosylated in the native polysaccharide (Fig. 3B). Next, we identified the sugars in these positions by re-examining the NMR spectra of MXP-A. From the linkage data, we knew that the missing residues were terminal Rha, terminal Glc, and terminal GlcNAc, as well as 4-linked Rha. Although we could not identify all the monosaccharide residues present in the NMR spectra of MXP-A (Fig. S5), we were able to partially assign most of the residues that were needed to complete the structure elucidation.

**Figure 3.**
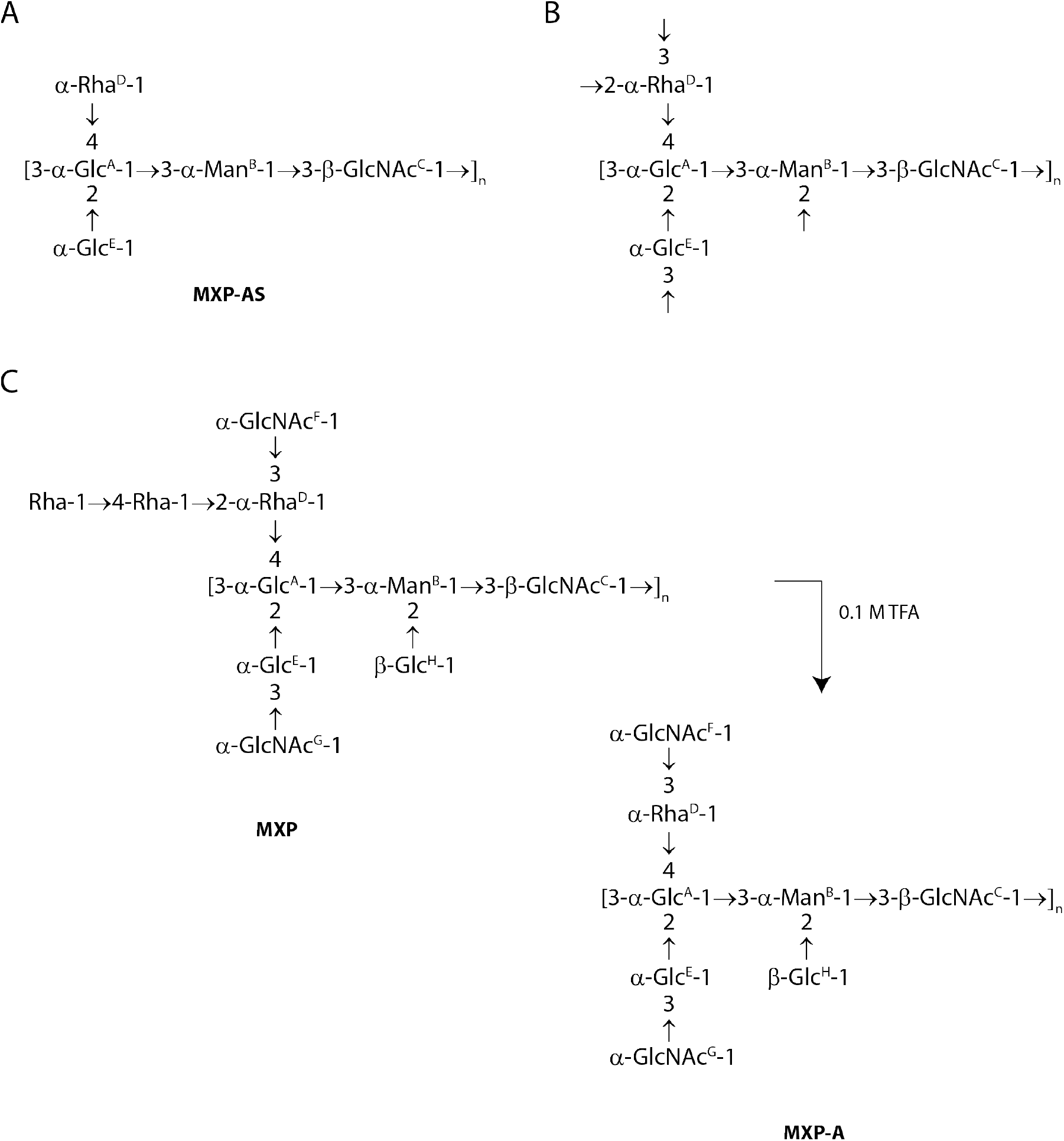
Elucidation of the chemical structure of the *Myxococcus xanthus* periplasmic high-MW carbohydrate (MXP). (*A*) Proposed chemical structure of the pentasaccharide repeat unit of the Smith-degraded MXP (MXP-AS). MXP-AS represents the core structure of the MXP; (*B*) linkages to the core structure MXP-AS that are cleaved during Smith degradation, based upon the linkage analysis results; (*C*) proposed structure of the decasaccharide repeat unit of the MXP and its product after mild acid hydrolysis with 0.1 M TFA (MXP-A).

It was beneficial for this process that the linkages involved mostly O-2 and O-3, i.e. positions that are relatively close to the anomeric center, so that only some of the protons and carbons in each residue had to be assigned in order to establish connectivities. Table S3 shows the assignments that were thus obtained. The 2,3,4-α-Glc residue was readily assigned because it had similar chemical shifts to its counterpart in MXP-AS. The chemical shifts of Man and Rha residues are very similar, in that both have downfield H-2 chemical shifts, and we found two residues with H-2 chemical shifts greater than 4 ppm. However, only one of the H-2 protons was correlated in the TOCSY spectrum with the H-6 methyl group of Rha, thus the other residue was identified as Man. Both of these residues were in the α-configuration, as evidenced by the downfield chemical shift of their anomeric protons. The Man residue had downfield chemical shifts for C-2 and C-3, showing glycosylation on O-2 and O-3. Although from the linkage analysis we expected to find 2,3-Rha in the NMR spectra, the Rha residue showed only a downfield displacement of C-3, whereas C-2 was more typical of an unsubstituted position. None of the other spin systems in the spectra of MXP-A were consistent with 2,3-α- or 2,3-β-Rha. Therefore, we concluded that the substituent that had originally been attached to O-2 of this Rha residue was lost during the initial mild acid hydrolysis. We were further able to identify and partially assign two terminal α-GlcNAc residues, a 3-linked β-GlcNAc residue, a 3-linked α-Glc residue, and a terminal β-Glc residue. NOE and HMBC correlations (Fig. S5) clearly indicated linkage of the β-Glc residue to O-2 of α-Man, establishing the location of this terminal residue. An NOE contact proved that the attachment site of the terminal α-GlcNAc residue **G** was on O-3 of Residue **E**. The connectivity of the other terminal α-GlcNAc (Residue **F**) was more difficult to discern because its anomeric proton was correlated to a signal that could belong to either H/C-3 of 3-α-Rha or 3-β-GlcNAc, as these two signals resonated at the same position in both proton and carbon dimensions. However, the α-GlcNAc residue could not be attached to the 3-β-GlcNAc, because Smith degradation would have liberated a terminal GlcNAc, but none was found in the linkage analysis of MXP-AS. Hence, the α-GlcNAc **F** is attached to O-3 of 3-α-Rha (Residue **D**). The location of the terminal and 4-linked Rha residues was determined by their absence in the NMR spectra. None of the spin systems detected in the spectra were consistent with either of these linkages in α- or β-form. Apparently these two residues constituted the side chain that was attached to O-2 of Residue **D** in MXP, but which was lost during acid hydrolysis. Thus, we conclude that a Rha-1-4-Rha disaccharide side chain is attached to O-2 of 2,3-α-Rha in MXP before acid hydrolysis. Combining all these results, the proposed structures of MXP and MXP-A are pictured in Fig. 3C.

### The synthesis of the periplasmic polymer is osmoregulated

Low-MW OPGs or OPG-like oligosaccharides are the only currently described periplasmic carbohydrates of Gram-negative bacteria; in virtually all cases their synthesis is osmoregulated (Lee et al., 2009; Bontemps-Gallo and Lacroix, 2015). Therefore, we next examined the osmoregulation of the periplasmic polymer. Since *M. xanthus* is grown in CTT medium (85 mOsm), our initial isolations demonstrated that the polymer was abundantly synthesized at low osmolarity. However, testing the high osmolarity scenario using liquid media was challenging, as the cells both do not tolerate the addition of salt and are strictly aerobic, and therefore sensitive to changes in oxygen availability caused by increasing viscosity through the addition of outer membrane-impermeable polymers such as polyethylene glycol (PEG). Therefore, we performed these experiments using cells grown on trays of solid CTT medium containing various concentrations of PEG (Chang et al., 2007) and gelrite as the solidifying agent, as agar does not solidify in the presence of PEG (Shungu et al., 1983). After confirming that replacing agar with gelrite had no effect on the synthesis of the polymer at low osmolarity, we repeated our isolations from cells grown at intermediate, 8% (0.5 MPa) and high, 16% (1 MPa) PEG concentrations (Chang et al., 2007). After harvesting the cells, we monitored the abundance of the polymer by examining both the crude fraction containing the polymer (Fig. 4A) and the purified 400 kDa large fragment (Fig. 4B). We were able to recover less of the periplasmic polymer at intermediate osmolarity (5% at 8% PEG) and none whatsoever at high osmolarity (16% PEG; Fig. 4). Of note, these data demonstrate that the polymer, although chemically very different from OPGs, physiologically phenocopies them including the osmoregulation of their synthesis.

**Fig. 4.**
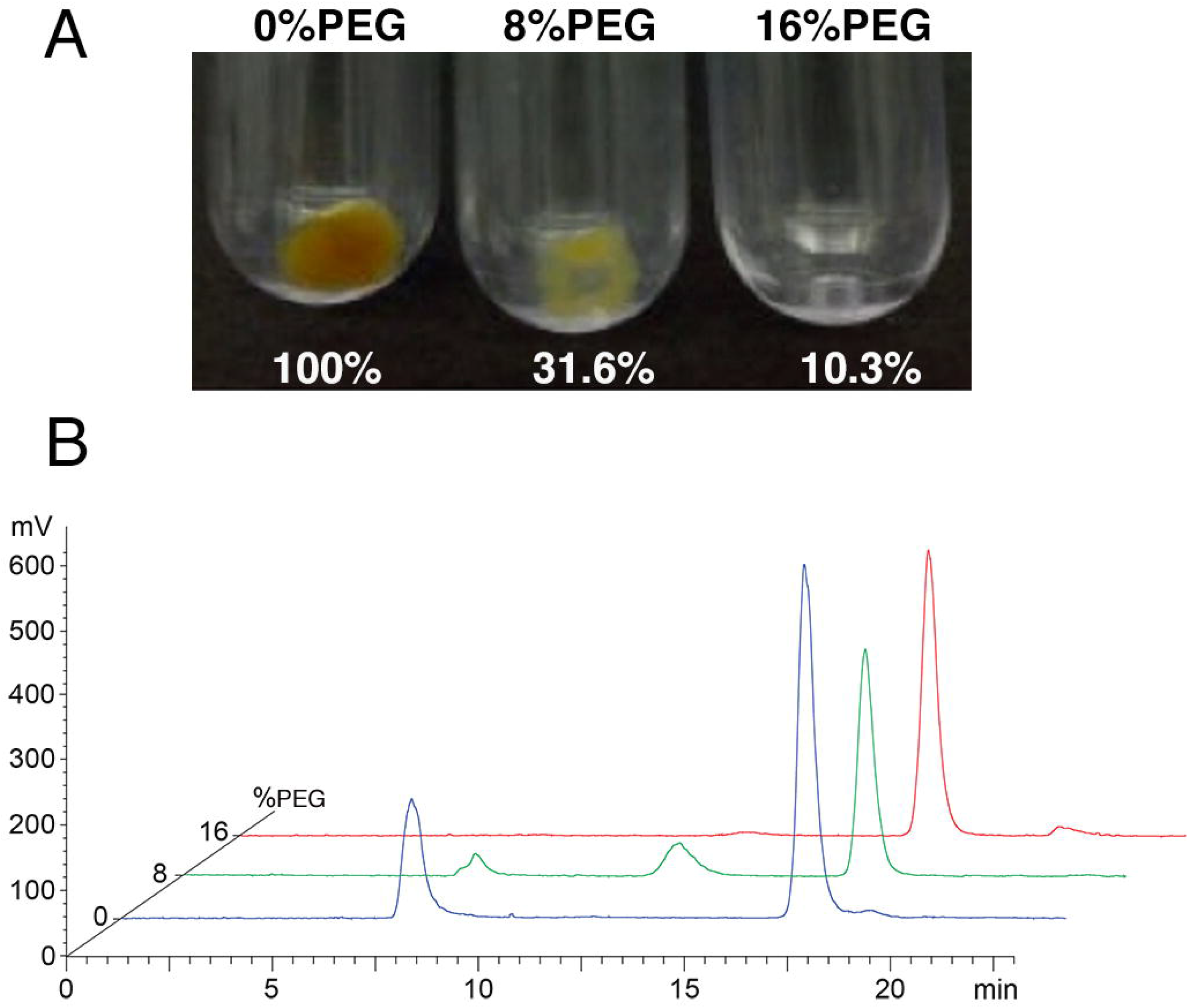
The *Myxococcus xanthus* high-MW carbohydrate is only produced by cells grown at low osmolarity. (*A*) Comparison of the amount of isolated crude polymer of cells grown under different osmotic conditions. Note that upon further purification no polymer can be recovered from the cells grown with 16% PEG. (*B*) Size exclusion chromatography of isolated polymer after extraction from cells grown under different osmotic conditions. The 400 kDa polymer peak decreases under intermediate (8% PEG) and disappears at high osmolarity (16% PEG). The peak that is present in all three isolations towards the end of the column runs does not contain carbohydrate.

### *M. xanthus* encodes OpgG and OpgH homologs

Although we were unable to isolate a canonical OPG, we searched the genome of *M. xanthus* for the presence of genes potentially involved in the synthesis of periplasmic glucans. Homologs of two core genes necessary for canonical OPG synthesis in most bacteria, *opgG* (*mxan_2385*) and *opgH* (*mxan_2383*) (Bontemps-Gallo et al., 2017), are present in *M. xanthus*. The primary amino acid sequences of OpgG and H in various species were compared. Whereas OpgG is highly conserved throughout its entire sequence among various bacteria (Fig. S1), OpgH sequences show a higher degree of variation, particularly at their N- and C-termini (Fig. S2). In *M. xanthus*, for example, the N-terminus is 124 aa shorter than in *E. coli*. We next attempted to delete these genes to test whether they are involved in the synthesis of the periplasmic high-molecular-weight carbohydrate. Repeated attempts at deleting *opgG* failed, indicating that this gene may be essential in *M. xanthus* grown under standard conditions. Efforts to delete *opgH* were similarly unsuccessful. As we were unable to delete these genes, and as no simple correlation between OPG protein sequences and synthesized polymers has been identified (Bontemps-Gallo et al., 2017), we are currently unable to assess whether OpgG and H in *M. xanthus* are involved in the synthesis of the high-molecular-weight periplasmic carbohydrate.

### The osmoregulated periplasmic polymer appears to protect and stabilize cells during the rod-to-sphere conversion of glycerol spore formation

In addition to environmental changes in osmolarity, vegetative cells of *M. xanthus* degrade their peptidoglycan during rod-to-spore morphogenesis (Bui et al., 2009; Higgs et al., 2014) and become highly sensitive to peptidoglycan-targeting antibiotics (White et al., 1968; Jones et al., 1981). Consistent with the osmoregulation of MXP we observed and known osmoprotection conferred by OPGs, we hypothesized that the fiber-forming periplasmic polymer may also function to mechanically stabilize the cells during the cell wall remodeling process during sporulation. To test this hypothesis, we grew vegetative cells in conditions under which they produce either ample (low osmolarity CTT) or virtually no polymer (high osmolarity CTT+16% PEG) and tested the cell’s ability to successfully form spores upon glycerol induction (Dworkin and Gibson, 1964). Although we cannot completely rule out that the different growth regimens affect other processes that influence spore formation, the results indicated that cells possessing the periplasmic polymer completely converted to spores within 2 h, while never more than ca. 40% of the cells lacking the polymer converted after 4 h or longer of stimulation, moreover, many of these cells failed to become completely spherical (Fig. 5). Therefore, polymer-lacking cells appear to be highly impaired in their ability to form spores both with respect of the speed as well as the completeness of the conversion. In summary, these data support the hypotheses that the fiber-forming polymer is important for osmoprotection, and moreover forms a periplasmic fibrillar meshwork that mechanically supports the cell envelope during rod-to-spore morphogenesis. In the latter case, it would support the mechanical robustness of the cell during the time before the newly synthesized spore coat replaces the degraded peptidoglycan layer, meaning that a complete conversion process would only be possible in the presence of the polymer (Bui et al., 2009; Higgs et al., 2014).

**Fig. 5.**
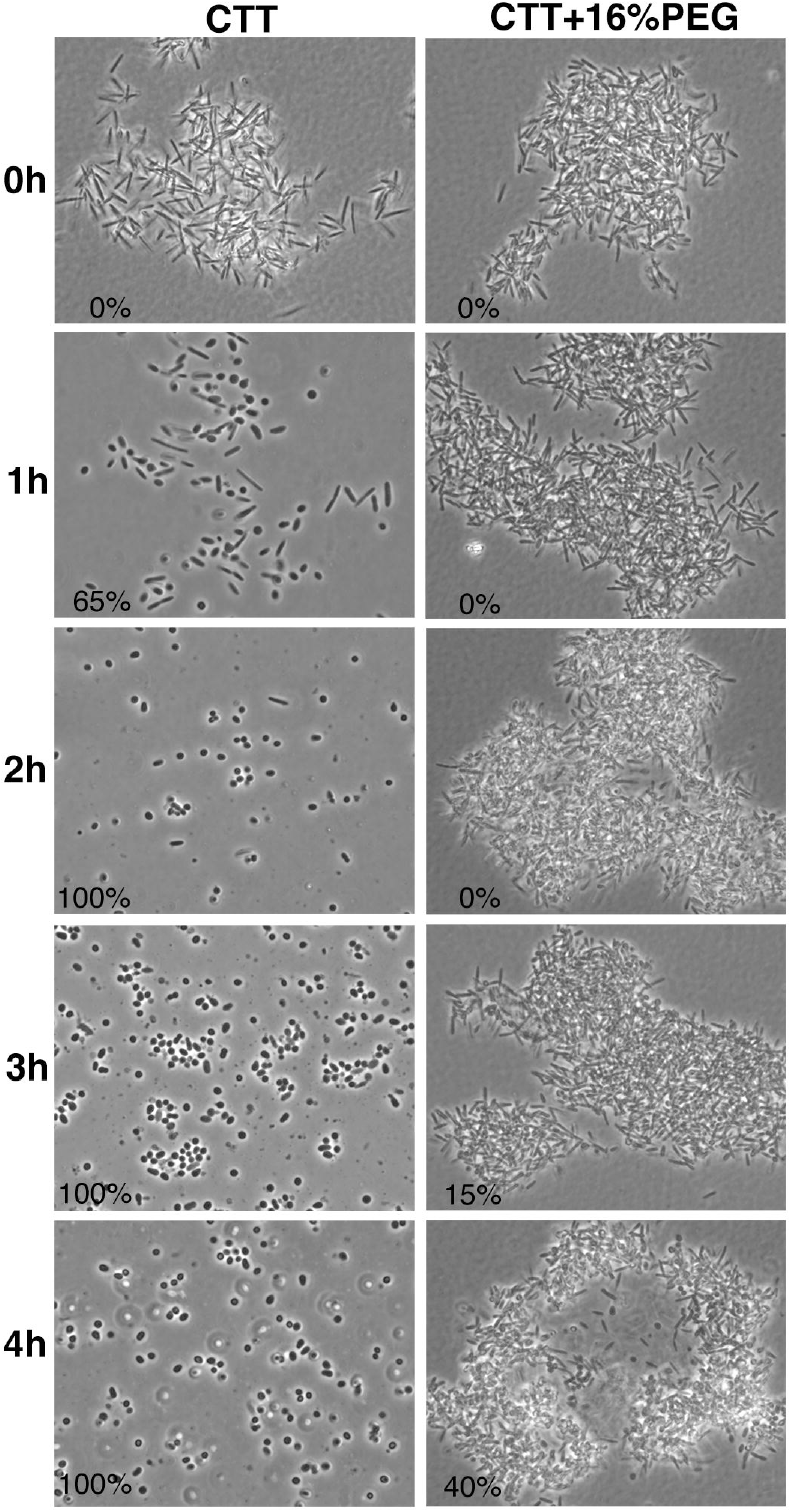
The presence of the high-MW polymer is correlated with the ability for cells to convert from rods to spores upon glycerol-induced sporulation. The column on the left shows wild type cells grown under low osmolarity in CTT medium. Under these conditions the cells contain large amounts of the polymer and nearly 100% convert to spores within ca. 2 h following induction. The cells on right were grown at high osmolarity in the presence of 16% PEG. These cells completely lack the periplasmic polymer and convert to glycerol spores substantially slower and to a greatly-reduced extent. Moreover, many of the converting cells do not complete the conversion but form shortened egg- or loaf-shaped cells instead. The small numbers in the pictures denote the percentage of cells converted to spores at each time point.

## Discussion

OPGs or OPG-like oligosaccharides are linear or cyclic 5-40 glucose molecule-long compounds that have been found in the periplasms of α-, β-, γ-, and ε-proteobacteria and that are an important part of the cell’s response to changes in environmental osmolarity (Bontemps-Gallo and Lacroix, 2015; Bontemps-Gallo at al., 2017). Here, we show that *M. xanthus,* a δ-proteobacterium, appears to lack a canonical OPG. Instead, the bacterium synthesizes a high-MW fiber-forming periplasmic carbohydrate that is chemically distinct from any known OPGs. First, it is substantially larger than canonical OPGs. Although we were not able to determine its native weight, we reason that it must be several thousand kilodaltons because the non-hydrolyzed polymer forms microscopically-visible, micrometer-long filaments and limited acid hydrolysis released a fragment of about 400 kDa. This is in strong contrast to the MW of canonical OPGs, which are usually smaller than 8 kDa (Bohin, 2000; Lee et al., 2009). Secondly, while all OPGs are polyglucans, this polymer contains three additional monosaccharides beside glucose, namely mannose, N-acetylglucosamine, and rhamnose. In addition, its decasaccharide repeat unit contains mostly α-glycosidic bonds, which differs from the β-glycosidic bonds dominating OPGs. Finally, OPGs are often modified with negatively charged groups resulting in a strong anionic character of these molecules. In contrast, no such substituents were detected in the polymer despite extensive analyses, indicating that this molecule appears to be neutral under all tested growth conditions. The only two aspects in which the polymer resembles canonical OPGs is its periplasmic location and the fact that its synthesis is osmoregulated, indicating that it may physiologically function like OPG. Intriguingly, bioinformatics identified in *M. xanthus* two core genes, *opgG* and *opgH* responsible for the synthesis of canonical OPGs in other bacteria. Repeated attempts to delete these genes failed, indicating that they appear to be essential under normal growth conditions. Therefore, we were unable to directly test their involvement in osmotolerance or MXP synthesis. In these other bacteria, no sequence-based correlation between *opg* genes and the chemistry of their products has been found, therefore we are unable to conclude that they do synthesize MXP. Consequentially, two scenarios could be possible: either *opgG* and *opgH* are involved in the synthesis of the high-MW periplasmic polymer, further indicating that correlation between *opg* genes and their products is virtually impossible, or the bacterium produces a canonical OPG that we were unable to detect. We do not currently have experimental evidence to support either of these hypotheses. However, based on the fact that the chemical composition and structure of MXP is so different from canonical OPGs, we consider it more likely that not yet identified glysosyltransferases are responsible for the synthesis of this polymer.

Together these observations strongly suggest that one of the main functions of the polymer is osmoprotection through the facilitation of a hydrated periplasmic space. As has been proposed for OPGs, this does not exclude the possibility that the polymer could also play other non-osmoregulated roles such as providing a cellular pool of monosaccharides, controlling periplasmic crowding, interacting with enzymes, influencing the solubility of compounds, or performing structural roles (Lee et al., 2009; Bontemps-Gallo et al., 2017). Because all of these functions can be accomplished by canonical OPGs, it raises the question why the periplasmic carbohydrate of *M. xanthus* is so different. Since no other δ-proteobacterium has so far been characterized with respect to its OPG, it is conceivable that δ-proteobacteria, or more specifically, myxobacteria, generally possess larger, more structurally complex OPGs. This could be explained by the evolutionary history, as the branch point between the δ-proteobacteria and the remaining proteobacteria is ancient, suggesting a distant genetic relationship (Emerson et al., 2007). However, we consider it more likely that this polymer in *M. xanthus* fulfills a function that is linked to the particular life cycle of this organism. *M. xanthus*, like other myxobacteria, is characterized by its unique ability to undergo developmental differentiation, a process during which starving cells coalesce into fruiting bodies, and eventually differentiate into spores (Zusman et al., 2007). While spore formation exists in other bacteria, the process in *M. xanthus* differs fundamentally from the analogous event in endospore-forming bacteria such as *Bacillus subtilis* (McKenney et al., 2013) in that the entire vegetative cell converts into a spherical spore without an initial asymmetric septation event (Higgins and Dworkin, 2012; Higgs et al., 2014). During this conversion process, the entire peptidoglycan of the vegetative cell is actively rearranged and finally completely degraded (Bui et al., 2009) while the cell simultaneously forms a spore coat that functionally replaces the peptidoglycan layer (Higgs et al., 2014; Holkenbrink et al., 2014). Consequently, the cells are particularly vulnerable to peptidoglycan-targeting antibiotics or to mechanical or osmotic stress during this rod-to-spore morphogenesis (White et al., 1968; Jones et al., 1981). As our experiments suggest, cells that possess the high-MW polymer can convert rapidly and completely (100% within 2 h) while those lacking the molecule are highly impaired in their ability to convert (with not more than 40% undergoing morphogenesis even after 4 h). The simplest explanation of this observation is that the fibrils of the polymer form a periplasmic meshwork that mechanically helps stabilize the cell envelope during the re-organization. This idea points to a highly specific function in *M. xanthus* suggesting that such large polymers may only be found in other spore-forming myxobacteria. However, since such polymers have not been identified in other species this interpretation is speculative; indeed, since the standard OPG isolation procedure using TCA does not release the polymer from *M. xanthus*, similar high-MW polymers may be produced by other bacteria but have been overlooked. For example, two studied bacteria, *Brucella* Sp. and *Sinorhizobium meliloti* strain GR4, possess OPGs that are not osmoregulated (Miller and Wood, 1996), which may point to the possibility that they possess other periplasmic molecules that are osmoregulated and involved in osmoprotection. Moreover, since high-MW carbohydrates are usually *per* definition classified as EPS, it may be that such polymers have already been discovered but misinterpreted as extracellular molecules. Ironically, such misidentification would mirror the initial discovery of OPGs, which were isolated from culture supernatants of *A. tumefaciens* and identified as crown gall polysaccharides and initially thought to be a specific sub-fraction of EPS (McIntire et al., 1942), before later studies revealed their periplasmic nature (Van Golde et al., 1973; Schulman and Kennedy, 1979). The discovery of the periplasmic polymer raises a new question, namely, the fate of the polymer after the conversion of the rods into spores. It will need to be resolved whether the polymer is kept in spores as part of the multilayered spore coat (Higgs et al., 2014; Holkenbrink et al., 2014) or whether it is enzymatically degraded and potentially used to synthesize spore coat material.

While the roles of the newly discovered periplasmic polymer in osmoregulation and cell envelope stabilization appear apparent, it likely influences additional cell envelope-related processes. One such process is A-motility in *M. xanthus*, a form of surface-associated locomotion that relies on large-scale interaction between the bacterial cell surface and the underlying substrate (Nan et al., 2014; Islam and Mignot, 2015). Although the precise molecular mechanism for this motility is still debated, it is clear that the cell envelope plays a major role. According to one model, the focal adhesion model (Mignot et al., 2007), motility is driven by the movement of motor complexes that reside at the cytoplasmic membrane and move along cytoskeletal tracks. As these motor complexes simultaneously bind to the extracellular substrate, they push the cells forward while moving backwards on the tracks (Faure et al., 2016). It has been noted that the presence of the peptidoglycan sacculus would potentially hinder the movements of such trans-envelope complexes (Nan et al., 2014); we suggest that the fibrillary periplasmic meshwork reported here may be an additional obstacle. In addition, this meshwork may influence important aspects of another A-motility model, the helical rotor model (Nan et al., 2010; Nan, 2017). Here, cytoplasmic cargo protein complexes are proposed to move backwards on a helically arranged cytoskeleton generating trans-envelope wave-like deformations that push the cell forward. An important constraint of this model is the ability of these protein complexes to generate waves with an amplitude large enough to be effective outside the cell. While it is difficult to judge how the periplasmic polymer would influence the transmission of such waves, the contribution of such structural feature to the envelope must be considered.

Another process to which the polymer could contribute is cellular flexibility. Although myxobacteria are physiologically like any other Gram-negative bacteria, they are extremely flexible, a characteristic they share i.e. with flexibacteria (Burchard, 1981; 1982). In contrast to i.e. *E. coli*, this allows *M. xanthus* to bend, turn, and twist, behaviors that have been exploited to examine their motility (Wolgemuth, 2005). No plausible ultrastructural explanation has been presented as to how the cells are able to bend at such great angles without snapping and breaking. The conventional view holds that the peptidoglycan is the mechanical structure that shapes the cells. However, since the periplasmic fibrillar meshwork certainly influences the overall mechanical properties of the cell envelope, we consider that this gel-like meshwork may behave like a hydroskeleton that can redistribute within the periplasm to accommodate strong physical deformations, or by doing so initiates rapid flexing of the cell body.

In addition, the meshwork could be involved in the strong tendency of *M. xanthus* cells to extrude outer membrane vesicles (Palsdottir et al., 2009). Although vesicle formation has been observed for every Gram-negative bacterial species (Kulp and Kuehn, 2010), myxobacteria stand out for the sheer amount of these structures often resulting in the formation of vesicle chains or tubes (Palsdottir et al., 2009; Berleman and Auer, 2013; Wei et al., 2014). Vesicle formation together with membrane fusions are at the basis of many aspects of myxobacterial multicellularity such as surface protein and LPS exchange (Pathak et al., 2012, Ducret et al., 2013; Wei et al., 2014; Vasallo et al., 2015), cell-to-cell signaling (Stevens and Søgaard-Andersen, 2005) and kin selection (Velicer and Vos, 2009; Vasallo et al., 2017). The presence of an osmotically active periplasmic carbohydrate could conceivably play a role in generating some of the force necessary to generate these important surface structures.

Finally, the periplasmic fibrillar OPG meshwork could certainly influence cell division as it is likely that the cells have to depolymerize the fibrils locally in order to allow septation and cell division to occur. This interpretation is based on the observation that any extended physical structure such as DNA or protein filaments needs to be either actively segregated or locally disassembled before cell division can occur (Badrinarayanan et al., 2015; Cabeen et al., 2009). Conversely, if cells lack the necessary clearing mechanisms, for example if foreign filament-forming proteins are overexpressed in *E. coli*, cell chaining often occurs as a consequence of incomplete divisions (Bharat et al., 2015). Of note, recent research has revealed a possible direct link between the synthesis of OPGs and cell size control and division in *E. coli* (Hill et al., 2013). Here one of the OPG-synthesizing enzymes, the glucosyltransferase OpgH interacts with the septal ring-forming protein FtsZ and, through sequestration, influences cell size and division. Through binding of UDP-glucose, OpgH measures nutrient availability and relates this information directly to the cell division machinery. Although this exact scenario is unlikely to be found in *M. xanthus*, the presence of a periplasmic polymer that forms micrometer-long fibrils indicates that the cells must either disassemble or segregate these structures within the periplasm for cell divisions to occur.

The discovery of this unique periplasmic OPG in *M. xanthus* identifies another major cellular component in this important model organism. It will be informative to determine whether other myxobacteria, δ-proteobacteria or bacteria in general, produce similarly complex OPGs. To a large extent this may depend on the principal function of this novel type of OPG. Future studies will be needed to clarify the identity of the enzymes responsible for the synthesis and assembly of the polymer, and examinations of mutants that fail to make the polymer will be needed to address the specific contributions of this polymer to cellular functions. Does the polymer address a specific need of the cell linked to the complex behavior and physiology of *M. xanthus*, does it allow adaptation to changing osmotic conditions that may be experienced by many environmental bacteria, or does it contribute to both?

## Materials and Methods

### Bacterial Strains and Growth Conditions

The *M. xanthus* strains used in this study were the wild-type strain (DK1622; Kaiser, 1979) and the EPS-deficient ΔdifE strain (YZ603; Black and Yang, 2004). Both strains were routinely grown in CTT or TPM medium with or without agar (Hodgkin and Kaiser, 1977). To test the influence of water limitation on the production of the periplasmic carbohydrate, either no, 8 or 16% of PEG 8000 (Fluka) were added to the CTT medium, which then was solidified using 1% gelrite (Shungu et al., 1983).

### Isolation and Purification of the Osmoregulated Periplasmic Carbohydrate

To isolate the osmoregulated periplasmic carbohydrate, wild type or ΔdifE cells were grown to mid-log phase in 2 × 400 ml CTT medium-containing flasks. This pre-culture was then either expanded into 8 × 400 ml flasks for liquid culture or harvested in a sterile centrifuge bottle, re-suspended in a small amount of medium and plated onto trays (30 ×40 cm) containing CTT agar (or gelrite) for solid surface growth. Both liquid and agar cultures were grown over night at 32 °C and harvested either by centrifugation or by scraping. The cell pellets were re-suspended in 120 ml extraction buffer (10 mM Tris-HCl pH 8.0, 10 mM MgSO_4_), transferred into a household blender, and blended for 3 min at high speed. The cell suspension was centrifuged twice to remove cells (10 min 27,500 × g, 10 min 47,800 × g) before 20% (w/vol) ammonium sulfate was added, and the precipitated polymer was collected (15 min at 47,800 × g). The pelleted polymer was re-suspended in water and extracted with 0.05% (w/vol) n-dodecyl β–D-maltoside to solubilize contaminating membranes. After centrifugation and re-suspension, the polymer was precipitated with (NH_4_)_2_SO_4_, centrifuged, and either dissolved in water or kept as a pellet and used for all analyses. To compare the extraction efficiency of various treatments, the cells were shaken by hand for 1 min, washed in 250 or 500 mM NaCl or treated by osmotic shock. After purification, the amount of extracted polymer was weighed and the variance determined by one-way ANOVA, while the p-values were calculated using the Tukey test.

### Isolation of Canonical Periplasmic Glucans

Canonical osmoregulated periplasmic glucans were extracted according to standard procedures (Cogez et al., 2001). Briefly, exponentially growing bacteria were harvested by centrifugation (10 min 27,500 × g) and the pellets were extracted with 5% trichloroacetic acid (TCA). After removal of the extracted cells (10 min 27,500 × g), the resulting supernatant was cleared of cell debris (10 min 47,800 × g), neutralized with ammonium hydroxide, desalted using a Sephadex G-15 column, and monosaccharides were analyzed after hydrolysis using 2 M TFA for 2 h at 121 °C.

### Osmotic Shock of Cells

Mid-log grown wild-type cells were harvested by centrifugation (10 min 27,500 × g). The cells were washed with ice-cold TM buffer (10 mM Tris-HCl pH 8.0, 8 mM MgSO_4_; 40 ml per gram cell pellet) and the wash supernatant was kept (Tris/MgSO_4_ wash). The cells were centrifuged again, re-suspended in the buffer and 20% sucrose was added. After 30 min on ice, the cells were centrifuged and the supernatant kept (sucrose wash). The cell pellet was rapidly dispersed in ice-cold TM, centrifuged, and the supernatant containing the periplasmic fraction kept (periplasm). Finally, the pelleted cells were re-suspended in 10 mM Tris pH 7.6 buffer containing cOmplete EDTA-free protease inhibitor cocktail (Roche Applied Sciences, Penzberg, Germany), sonicated, and centrifuged to isolate the cytoplasmic fraction (cell pellet). Both, colorimetric assays such as the anthrone test and biochemical isolations using the described protocol indicated that only the periplasmic fraction contained the carbohydrate polymer.

### Alkaline Phosphatase Assay

The SensoLyte pNPP Alkaline Phosphatase Assay Kit (AnaSpec, Fremont, CA) was used according to the manufacturer’s instruction to measure the activity of the enzyme in each supernatant fraction obtained from the osmotic shock procedure (Tris/MgSO_4_ wash, sucrose wash, periplasm, and cell pellet). Briefly, a 96 well plate was used, and 50 μl of supernatant samples or phosphatase standard solutions were pipetted into the wells. Alkaline phosphatase activity was measured by adding 50 μl of p-nitrophenyl phosphate phosphatase (pNPP) substrate solution to each of the wells. The plate was incubated for 60 min at room temperature, shaken for 1 min, and the absorbance measured at 405 nm in a plate reader (BioTek Industries, Winooski, VT).

### Antibody Production

Isolated purified periplasmic carbohydrate was injected into rabbits (Cocalico, Reamstown, PA) to generate polyclonal antibodies according to standard protocol. Both the test bleeds and the collected sera were tested for cross-reactivity using a dot blot assay and various mono- and polysaccharides. The membranes were treated according to standard protocols and visualized using the Pico chemiluminescent reagent (Pierce, Rockford, IL).

### Osmolality Measurements

To determine the osmolality of solutions a Micro-Sample Osmometer Model M3 (Advanced Instruments, Norwood, MA) was used. About 20 μl of the samples was drawn into the capillary of the sample holder and measured using freezing point depression. The instrument was calibrated using standard solutions of low, medium, and high osmolality.

### Anthrone Test

Five milliliters of freshly prepared ice-cold anthrone reagent (0.2% in 95% H_2_SO_4_) were added to 1 ml of sample or variously diluted glucose standard solutions (10 mg/ml). Samples were mixed, heated for 10 min, rapidly cooled, and the absorption at OD_620_ determined (Seifter et al., 1950).

### Analytical Carbohydrate Methods

#### Dialysis

To remove salts and low-molecular-weight contaminants, the purified carbohydrate was dialyzed for 48 h against running de-ionized water using a Slide-A-Lyzer dialysis cassette with a molecular cutoff of 20 kDa (Pierce, Rockford, IL). Dialyzed MXP samples were freeze-dried and stored at −20 °C for further analysis.

#### Partial Hydrolysis

Five milliliters of 0.1 M trifluoroacetic acid (TFA) were added to 20 mg freeze-dried polysaccharide, and the mixture was incubated for 2.5 h at 80 °C. After partial hydrolysis, methanol was added the liquid was evaporated under a dry stream of N_2_ to remove the acid. The dried samples were dissolved in de-ionized water and filtered through a 0.22 μm filter to give about 17 mg MXP-A.

#### Size Exclusion High-pressure Liquid Chromatography (SEC-HPLC)

For carbohydrate fractionation, a Superose 12 gel filtration column (GE Healthcare Life Sciences, Little Chalfont, UK) was used. A total of about 200 μg partially hydrolyzed polysaccharide was loaded onto the column at a concentration of ca. 1 mg/ml and eluted at a flow rate of 0.5 ml/min using 50 mM ammonium acetate buffer pH 5.2. Eluting carbohydrates were detected with an ELS detector (evaporation temperature 70 °C, gain 9, filter 5) and data were collected and processed using the Agilent ChemStation software (Agilent, Santa Clara, CA). For calibration, dextran standards with average molecular weights of 551, 167, 67, 40, and 5 kDa, as well as maltoheptaose, maltopentaose, and glucose were used.

#### Smith Degradation

Smith degradation was carried out as described previously (MacLean and Perry, 2010). The MXP-A (~4.5 mg) in water containing sodium metaperiodate (NaIO_4_, 9 mg/ 450 μL) was kept in the dark at 20 °C for 20 h. Following the addition of glycol (0.1 mL), the reaction mixture was dialyzed against 3 changes of 4 L deionized water. The retentate was treated with NaBH_4_ (10 mg/2 mL water, 4 h), followed by neutralization (AcOH) and further dialysis. The concentrated retentate was dissolved in 1 M hydrochloric acid (2 mL) and was kept at room temperature for 16 h, diluted with water, and the lyophilized product was fractionated by SEC column chromatography to yield several fractions.

#### Glycosyl Composition Analysis

Glycosyl composition analysis was performed by combined gas chromatography/mass spectrometry (GC-MS) of the per-*O*-trimethylsilyl (TMS) derivatives of the monosaccharide methyl glycosides generated by acidic methanolysis (Santander at al., 2013; Edgar et al., 2016). Briefly, sample aliquots were added into separate tubes containing 20 μg inositol as internal standard. After drying, methyl glycosides were prepared by mild acid treatment by methanolysis in 1 M HCl in methanol at 80 °C for 16 h, followed by re-*N*-acetylation with pyridine and acetic anhydride in methanol (for detection of amino sugars). Samples were then per-*O*-trimethylsilylated at 80 °C for 0.5 h using Tri-Sil (Pierce, Rockford, IL), separated on an Agilent DB-1 fused silica capillary column (30 m × 0.25 mm), and analyzed by GC-MS using an Agilent 7890A GC interfaced to an Agilent 5975C mass selective detector (MSD).

#### Glycosyl Linkage Analysis

For glycosyl linkage analysis, samples were permethylated, depolymerized, reduced, and acetylated, and the resulting partially methylated alditol acetates (PMAAs) analyzed by GC-MS as described previously (Heiss et al., 2009), but with slight modifications. Briefly, dry carbohydrate samples were suspended in 200 μl of dimethyl sulfoxide (DMSO) and continuously stirred for 1-2 weeks. The samples were then permethylated through treatment with sodium hydroxide and methyl iodide in dry DMSO. To ensure complete methylation of the polymer, the procedure was carried out in two steps. Initially, the polymer was subjected to NaOH for 15 min, before methyl iodide was added and left for 45 min. In the second step, additional base was added for 10 min, followed by more methyl iodide for another 40 min. The permethylated polymer was hydrolyzed using 2 M TFA for 2 h at 121 °C, reduced with NaBD_4_ and acetylated with a mixture of acetic anhydride and TFA. The resulting PMAAs were separated on a 30 m × 0.25 mm ID Supelco 2380 bonded phase fused silica capillary column (for neutral sugars) and on a 30 m × 0.25 mm ID Phenomenex ZB-1MS column (for amino sugars) and analyzed on an Agilent 7890A GC interfaced to a 5975C MSD using electron impact ionization mode.

#### NMR Spectroscopy

Polymer samples were deuterium exchanged by dissolving in D_2_O and lyophilizing. After deuterium-exchange, the sample was dissolved in 0.5 ml D_2_O and placed into a 5 mm NMR tube. 1-D proton and 2-D gCOSY, TOCSY, gHSQC, gHMBC, NOESY spectra were obtained on a Varian Inova 600 and 800 MHz spectrometer (Agilent Technologies, Palo Alto, CA) at 45 °C using standard Varian pulse sequences. Chemical shifts were measured relative to internal acetone (δ_H_=2.218, δ_C_=33.0 ppm).

#### Glycerol Spore Formation

To analyze rod-to-spore morphogenesis, cells were grown on trays with gelrite-solidified CTT medium with or without PEG, harvested, and re-suspended in 25 ml of CTT medium. Spore formation was induced by adding 0.5 M glycerol to the medium either immediately following the re-suspension or after an initial incubation of the cells for 1.5 h at 32 ºC. Samples were collected after shaking at 250 rpm for 0, 15, 30, 60, 90, 120, 180, and 240 min at 32 ºC, fixed and analyzed using the light microscope (Dworkin and Gibson, 1964). The assay was performed three times in triplicate. To determine the rate of conversion one hundred randomly selected cells were counted and the number of spherical spores *vs.* vegetative cells determined.

#### Electron and Light Microscopy

Negative staining was used to examine the structure of the isolated periplasmic polymer. Glow discharged-treated 400 mesh carbon-coated copper grids were placed on top of a drop containing water-dissolved polymer for 1-2 min and stained with either un-buffered 2% uranyl acetate or pH 7.5-buffered 2% phosphotungstic acid. For cryo-immuno electron microscopy, cells were processed using standard procedures (Tokuyasu, 1973). Sections were cut with a Leica UCT cryo-microtome (Leica Microsystems, Buffalo Grove, IL) and picked up on 200 mesh formvar-coated glow discharged nickel grids. For labeling, the grids were transferred onto drops of PBS containing the primary antibody at concentrations of 1:500, 1:1000, or 1:2000 and incubated overnight at 4 ºC. Detection of the primary antibody was done with 12 nm gold-labeled goat anti-rabbit secondary antibody (Jackson ImmunoResearch, West Grove, PA) diluted 1:20 in PBS for one hour at room temperature. Finally, the thin sections were contrasted using 2% methylcellulose containing 0.3% uranyl acetate for 10 min at 4 ºC. Samples were viewed with a Philips CM120 (FEI, Hillsboro, OR) or a Zeiss Libra (Zeiss, Thornhill, NJ) electron microscope operating at 80 kV and pictures were recorded using a 4×4 ORCA CCD camera (Hamamatsu, Middlesex, NJ). A Nikon E800 light microscope equipped with a 100× oil immersion phase contrast objective was used to take pictures of the glycerol spore formation (Nikon Instruments, Melville, NY). All recorded images were imported into Adobe Photoshop and digitally processed for publication. If the contrast was adjusted, it was applied to the entire image. No other form of digital alteration was performed.

#### Bioinformatics

OpgG and OpgH homologues in various bacteria were identified using standard BLAST searches (Altschul et al., 1990). The identified sequences were then used to generate a multiple sequence alignment using Clustal Omega (Madeira et al., 2019), which was visualized with Jalview2 (Waterhouse et al., 2009).

## ACKNOWLEDGMENTS

We would like to thank past and present members of the Hoiczyk laboratory for helpful discussions and comments on the work and the manuscript, Carol Cooke for help with the immuno-EM work, Mike Delannoy for assistance with osmolality measurements, Jane Voss for help with the photography, and Sara Porfirio for proof reading. This work was started at Johns Hopkins Bloomberg School of Public Health and then continued and finished at the University of Sheffield. The initial research was funded by a Sommer Scholar Fellowship (to S.A.), while completion was supported by funds of the Imagine initiative of the University of Sheffield (D.M.Z., K.S, J.M.T.S. and E.H.) and the BBSRC (E.H.). The work at the Complex Carbohydrate Center was supported by the Chemical Sciences, Geosciences and Biosciences Division, Office of Basic Energy Sciences, U.S. Department of Energy grant (DE-SC0015662 to P.A.).

