## Supplemental Material for "Structure and function of a novel osmoregulated periplasmic fiber-forming high-molecular-weight carbohydrate of *Myxococcus xanthus*"

### Supplementary Tables and Figures:

**Table S1:** Glycosyl linkage analysis of the polysaccharide before (MXP) and after acid hydrolysis and Smith degradation (MXP-AS).

| Linkage | MXP | MXP-AS |
|---|---|---|
| t-Rha | 2.4 | 5.5 |
| 4-Rha | 7.4 | 0.0 |
| t-Man | 0.0 | 1.2 |
| 3-Rha | 0.0 | 1.9 |
| t-Glc | 6.5 | 28.6 |
| 2,3-Rha | 6.7 | 0.4 |
| 3-Glc | 9.2 | 5.3 |
| 3-Man | 1.5 | 17.6 |
| 2-Glc | 0.0 | 1.4 |
| 6-Glc | 0.3 | 1.1 |
| 4-Glc | 0.2 | 2.2 |
| 2,3-Man | 10.9 | 0.9 |
| 3,4-Glc | 0.6 | 3.6 |
| 2,3-Glc | 0.3 | 7.2 |
| 2,3,4-Glc | 13.0 | 18.6 |
| 3,4,6-Glc | 0.0 | 1.6 |
| t-GlcNAc | 20.7 | 0.0 |
| 3-GlcNAc | 20.2 | 2.9 |

**Table S2.** Chemical shift assignments of MXP-AS.

| No. | Residue | | Chemical shift (ppm) | | | | | | NOE[a] |
|---|---|---|---|---|---|---|---|---|---|
| | | | 1 | 2 | 3 | 4 | 5 | 6 | HMBC[b] |
| A | 2,3,4-α-Glc | $^{1}H$ | 5.46 | 3.88 | 4.52 | 3.65 | 3.89 | 3.85/ND[c] | B3 |
| | | $^{13}C$ | 99.8 | 77.9 | 77.4 | 78.1 | 74.0 | 63.6 | B3 |
| B | 3-α-Man | $^{1}H$ | 5.26 | 4.14 | 3.87 | 3.88 | 3.58 | ND | C3 |
| | | $^{13}C$ | 103.4 | 72.9 | 80.9 | 68.6 | 76.0 | ND | C3 |
| C | 3-β-GlcNAc | $^{1}H$ | 4.87 | 3.89 | 3.74 | 3.60 | 3.47 | ND | - |
| | | $^{13}C$ | 102.6 | 57.4 | 82.6 | 73.6 | 78.7 | ND | A3 |
| D | α-Rha | $^{1}H$ | 5.14 | 4.24 | 3.73 | 3.47 | 3.69 | 1.28 | A4 |
| | | $^{13}C$ | 104.9 | 72.9 | 73.1 | 74.7 | 72.8 | 19.4 | A4 |
| E | α-Glc | $^{1}H$ | 5.22 | 3.59 | 3.72 | 3.47 | ND | ND | A2 |
| | | $^{13}C$ | 98.6 | 74.0 | 76.3 | 72.7 | ND | ND | A2 |

[a] inter-residue NOE correlations found between H-1 of the residue assigned in each row and the closest proton in the adjacent residue, designated by residue letter and proton number

[b] inter-residue HMB correlations found between H-1 of the residue assigned in each row and the linking carbon in the adjacent residue, designated by residue letter and proton number

[c] ND = not determined

**Table S3:** Chemical shift assignments of MXP-A.

| No. | Residue | Chemical shift (ppm) | | | | | | NOE[a] |
|---|---|---|---|---|---|---|---|---|
| | | 1 | 2 | 3 | 4 | 5 | 6 | HMBC[b] |
| A | 2,3,4-α-Glc | 5.39 | 3.89 | 4.46 | 3.60 | 4.05 | ND[c] | B3 |
| | | 99.5 | 75.4 | 76.3 | 78.3 | 73.8 | ND | - |
| B | 2,3-α-Man | 5.26 | 4.16 | 3.93 | 3.92 | 3.88 | ND | C3 |
| | | 101.5 | 79.4 | 77.9 | 68.6 | 76.2 | ND | C3 |
| C | 3-β-GlcNAc | 5.02 | 3.94 | 3.69 | 3.56 | ND | ND | A3 |
| | | 103.0 | 55.2 | 83.8 | 73.7 | ND | ND | - |
| D | 3-α-Rha | 5.05 | 4.32 | 3.70 | 3.58 | 4.03 | 1.27 | A4 |
| | | 105.9 | 72.0 | 83.1 | 73.7 | 73.9 | 19.6 | A4 |
| E | 3-α-Glc | 5.39 | 3.62 | 3.77 | ND | ND | ND | - |
| | | 96.4 | 72.1 | 81.2 | ND | ND | ND | - |
| F | α-GlcNAc | 5.11 | 3.89 | 3.77 | 3.74 | ND | ND | D3 |
| | | 101.8 | 56.9 | 73.9 | 73.2 | 72.7 | ND | D3 |
| G | α-GlcNAc | 5.51 | 4.00 | 3.70 | 3.62 | 3.98 | ND | E3 |
| | | 100.7 | 56.2 | 74.1 | 72.8 | 74.5 | ND | - |
| H | β-Glc | 4.35 | 3.28 | 3.45 | 3.53 | 3.32 | 3.86/3.74 | B2 |
| | | 104.0 | 75.4 | 78.5 | 72.1 | 78.8 | 63.5 | B2 |

[a] inter-residue NOE correlations found between H-1 of the residue assigned in each row and the closest proton in the adjacent residue, designated by residue letter and proton number

[b] inter-residue HMB correlations found between H-1 of the residue assigned in each row and the linking carbon in the adjacent residue, designated by residue letter and proton number

[c] ND = not determined

**Figure S1:** Multiple sequence alignment of OpgG sequences of *Myxococcus xanthus* (M.x., gene: *mxan_2385*, protein accession: ABF92764.1), *Escherichia coli* (E.c., gene: *opgG*, protein accession: AAC74132.1), *Pseudomonas syringae* (P.s., gene: ALP83_01217, protein accession: RMR57953.1), *Dickeya chrysanthemi* (D.c., gene: Dd1591_1604, protein accession: ACT06458.1), *Xanthomonas campestris* (X.c., gene: KWM_0108235, protein accession: KFA10717.1), *Ralstonia solanacearum* (Ra.s., gene: *mdoG*, protein accession: AKZ25463.1), *Rhodobacter sphaeroides* (Rh.s., gene: Rsph17029_1762, protein accession: ABN76872.1), and *Nitrosomonas europaea* (N.e., gene: *mdoG*, protein accession: CAD86349.1). The selected sequences are from bacteria representing various classes (Rh.s., alphaproteobacteria; Ra.s., N.e. betaproteobacteria; E.c., P.s., D.c., X.c. gammaproteobacteria; M.x. deltaproteobacteria) as well as various families of OPGs (class I: E.c., P.s., D.c; class IV: X.c., Ra.s., Rh.s.). Overall the percentage identities of the proteins range from 40% (N.e.) to 49% (Ra.s.) and 59% (N.e.) to 63% (Ra.s.) with MXAN_2385 from *M. xanthus*. Assignment of secondary structure elements is based on the X-ray structure of the *E. coli* protein (PDB entry 1TXK; Hanoulle et al., 2004), orange arrows denote beta sheets, while purple barrels denote alpha helices. The rectangle with the dashed outline at the N-terminus represents the signal sequence of OpgG of *E. coli*.

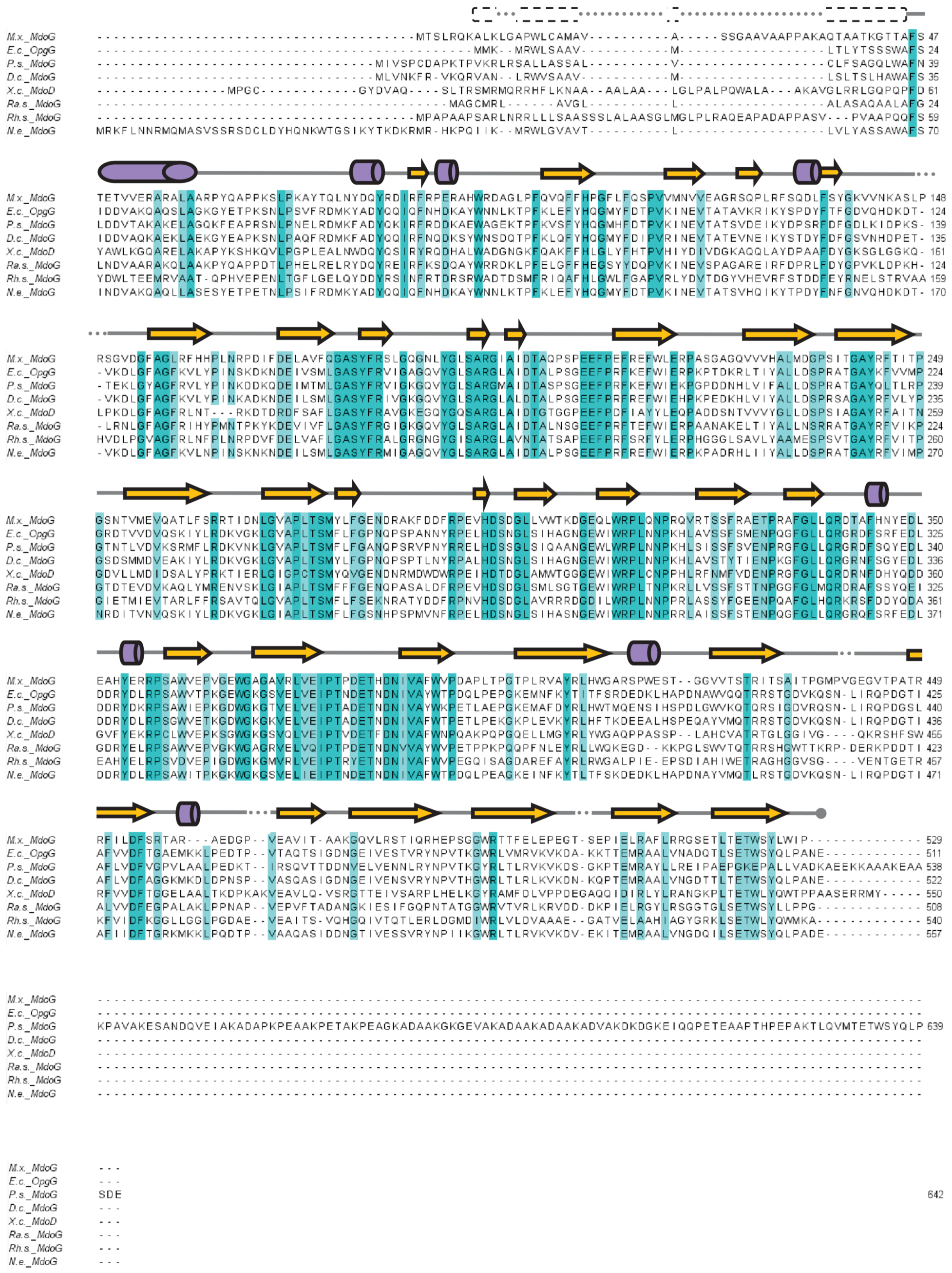
M.x._MdoG
E.c._OpgG
P.s._MdoG
D.c._MdoG
X.c._MdoD
Ra.s._MdoG
Rh.s._MdoG
N.e._MdoG

**Figure S2:** Multiple sequence alignment of OpgH sequences of *Myxococcus xanthus* (M.x., gene: *mxan_2383*, protein accession: ABF89910.1), *Escherichia coli* (E.c., gene: *opgG b1049*, protein accession: AAC74133.1), *Pseudomonas syringae* (P.s., gene: *ALP83_04833*, protein accession: RMR57981.1), *Dickeya chrysanthemi* (D.c., gene: *Dd1591_1605*, protein accession: ACT06459.1), *Xanthomonas campestris* (X.c., gene: *KWM_0110385*, protein accession: KFA09757.1), *Ralstonia solanacearum* (Ra.s., gene: *ACH51_03260*, protein accession: AKZ25464.1), *Rhodobacter sphaeroides* (Rh.s., gene: *Rsph17029_1764*, protein accession: ABN76874.1), and *Nitrosomonas europaea* (N.e., gene: *NE2438*, protein accession: CAD86350.1). The selected sequences are from bacteria representing various classes (Rh.s. alphaproteobacteria; Ra.s., N.e. betaproteobacteria; E.c., P.s., D.c., X.c. gammaproteobacteria; M.x. deltaproteobacteria) as well as various families of OPGs (class I: E.c., P.s., D.c; class IV: X.c., Ra.s., Rh.s.). Overall the percent identities and similarities of the proteins ranges from 39% (Rh.s.) to 47% (Ra.s.) and 55% (Rh.s.) to 62% (D.c.) when compared to MXAN_2383 of *M. xanthus*.

```
M.x._MdoH        ---------------------------------------------------------------------------------------------------
E.c._OpgH      1 ---------------------------MNKTTEYIDAMPIAASEKAALP------------KTDIRAVHQALDA-EHRTWA--REDDSPQGSVKARLEQ 57
P.s._MdoH      1 ----------------MSCRAMSNSLPVPVSLNEYLAHLPMSDEQRAELAG----------CKTFAELHERLSAQPV---N--DPAEAAQASVGRRLTL 68
D.c._MdoH      1 -------------------------MNKSTTSLEYIEKLPLPAGQAEALHEILS---S---SQDLSSVHQSLSDGSVVNMQ--SSDDIPLVSVPRRLEL 66
X.c._MdoH        ---------------------------------------------------------------------------------------------------
Ra.s._MdoH     1 MELPATSGLNAQPGNVEGTTASTRPATALSVAERYLEALPLPAEARAALRREAGIEPADKDAVALAKLHRALARLDAAQTASIEGSVPAYASIGSRLDA 99
Rh.s._MdoH       ---------------------------------------------------------------------------------------------------
N.e._MdoH      1 ---------------------------MNKTTEYIDALPLTVAEKATLP------------ATDIRTLHETLNP-EHHDYA--REDDSPLGSVKARLEQ 57

M.x._MdoH      1 -------------------------------------------------------------------------------------MHAHSFSPESAGIR 14
E.c._OpgH     58 AWPDS------LADG-QLIKDDEGRDQLKAMPEAKRSSMFPDPWRTNPVGRFWDRL----RGRDVTPRYLARLTKEEQES-------EQK-WRTVGTIR 137
P.s._MdoH     69 TTADQ------LEDAEMLGVDASGRLCLKATPPIRRTKVVPEPWRTNILVRGWRRL----TGKTNPPKPAHD-DLPRDLP-------KAR-WRTVGSIR 148
D.c._MdoH     67 AWEDG------LNGGKQLGKDREGRTALQAMPRITRASMFPDAWQTNPMARWWDSI----NGRTKPLRHQY-KTAEENEA-------ENR-WRKVGTLR 146
X.c._MdoH      1 ------------------------------MPPV----------------------SALDAGKPTLPPEAPLAMPEQNLREGSLQVRHQRTSPAGIGVR 47
Ra.s._MdoH   100 AYGTPHAGTATPEPAPPLEHDTAGRIHLDTGPEPARRSMVPWPWVLGPIWRARRAIGRLFSGGTAPVPY---EQPDSPDP-------KGI-WRFVGARR 187
Rh.s._MdoH     1 -------------------------------------------------------------------------------------MPAERRRRAVTLAS 14
N.e._MdoH     58 SWPDS------LVNR-QLTEDDEGRTQLETMPKATRSSISPDPWRTNPIGRFWDHL----RGHNATPHHVSRLTKEEQAH-------EQK-WCTVGTIR 137

M.x._MdoH     15 RLFVLGLAAVSTLVGTWEMHRLLSARGTTVPE----------------------GVLLVLFALCFGWIALSFWTAVAGFLQLAVSKRLPGLRWPTEEES 91
E.c._OpgH    138 RYILLILTLAQTVVATWYMKTILPYQGWALINPMDMVGQDLWVSFMQLLPYMLQTGILILFAVLFCWVSAGFWTALMGFLQLLIGRDKYSISASTVGDE 236
P.s._MdoH    149 RYILLILMLGQTIVAGWYMKGILPYQGWSLVSLDEITRQTFVQTALQVMPYALQTSILLLFGILFCWVSAGFWTALMGFLELLTGRDKYRISGASAGNE 247
D.c._MdoH    147 RYILLVLTLFQTAIATWYMKTILPYQGWALIDPFAMAHQDIWRTIMQLLPYVLQSGILILFAILFCWVSAGFWTALMGFLQLLIGKDKYSISSTITGNE 245
X.c._MdoH     48 RFYLIGGTFATTAVAVWVMLSVLWPDGISVLE----------------------GCLLGLFILLFAWIAMSFASAVAGFVTVVARAGRK--LGIDPEEP 122
Ra.s._MdoH   188 RLTLLALMIAQTVAATWAMSSVLPYHGHDWLE----------------------AIIIALFALLFCWVSAGFWTAITGFLLLAFHGDRFVISRRAAPNA 264
Rh.s._MdoH    15 RLVAAAISLTAAAGAFFLFLQFGSTDGLDSM---DIT-------------------RSVLILVSTSWLGWGAAHAVLGLFSR---PQRP-----ANVSP 83
N.e._MdoH    138 RYILLLLTFSQTALATWYMKTILPYQGWALIDPIDMIGQDIWISFMQLLPYILQSGILILFAILFCWISAGFWTALMGFLQLLIGKDKYSISASLAGDV 236

M.x._MdoH     92 ATRLTSRTSVVMPVHNEDPASVFANVQATYESVEATGQLDAFDFYILSDSTRPEAWIAEELAWADLCRRVGGQGRIFYRRRTDNTGKKAGNLQDFCERW 190
E.c._OpgH    237 PLNPEHRTALIMPICNEDVNRVFAGLRATWESVKATGNAKHFDVYILSDSYNPDICVAEQKAWMELIAEVGGEGQIFYRRRRRRVKRKSGNIDDFCRRW 335
P.s._MdoH    248 PIEKGARTALVMPICNEDVPRVFAGLRATFESVAATGDLDRFDFFVLSDTNETDIAVAEQQAWLDVCRETKGFGKIFYRRRRRRVKRKSGNLDDFCRRW 346
D.c._MdoH    246 PLNPKYRTALIMPICNEDVERVFAGLRATYESVAATGDLDHFDIYVLSDSNDPDICIAEQKAWMDLCREVDGAGRIFYRRRRRRVKRKSGNIDDFCRRW 344
X.c._MdoH    123 LPTLRSRTALLMPTYNEDPRRLLAGLQAIYESVAETGQLEHFDFFVLSDTTREHIGRAEELVYGELCDRVDGHGRIFYRRRADNAARKAGNVADWVRRF 221
Ra.s._MdoH   265 PIPDDARTAIVMPICNEDVQRVFAGLRATYESLQRTGHLEHFDFFVLSDSGDPDIRTAEADAWLHVCRALDGFGRVFYRWRRHRVKRKSGNIDDFCRRW 363
Rh.s._MdoH    84 DAPISTRTVILVPVYNEDPVATFSRIAAMDASLAATPWRDLFHFAILSDTRDEAIAARERFWFLRLLRERDAEGRIFYRRRAVNRGRKAGNIEDFIQKS 182
N.e._MdoH    237 PINPEHRTALIMPICNEDVDRVFAGLRATWESVKATNQQQHFDIYILSDSYDPDICVAEQKAWIELLAEVQDKGQIFYRRRGRRVKRKSGNIDDFCRRW 335

M.x._MdoH    191 GRHYDFTIVLDADSLMDGRTLVTMARLMELNPRAGILQAPPLNVGRSTLFARLQQFAGRVYGPVVAAGAAAWQLGESNYWGHNAIIRTEAFIQHCGLPV 289
E.c._OpgH    336 GSQYSYMVVLDADSVMTGDCLCGLVRLMEANPNAGIIQSSPKASGMDTLYARCQQFATRVYGPLFTAGLHFWQLGESHYWGHNAIIRVKPFIEHCALAP 434
P.s._MdoH    347 GGEYRYMVVLDADSVMSGECLTSLVRLMEATPDAGIIQTAPRASGMDTLYARMQQFATRVYGPLFTAGLHFWQLGESHYWGHNAIIRMKPFIEHCALAP 445
D.c._MdoH    345 GNQYSYMVVLDADSVMSGECLTSLVRLMEANPNAGIIQSAPRASGMDTLYARCQQFATRVYGPLFTAGLHFWQLGESHYWGHNAIIRVKPFIEHCALAP 443
X.c._MdoH    222 GGSYSQMLILDADSVMTGDTIVRLVAGMENNPDVGLIQTLPAVVNGQTLFARMQQFGGRVYGPIIAFGVAWWHGAESNYWGHNAIIRTQAFADHAGLPS 320
Ra.s._MdoH   364 GGRYRYMIVLDADSVMSGDCLTRLVQLMEGAPSAGIIQTAPLAAGRDTLYARIQQFATRVYGPLFTAGLHYWQLGESHYWGHNAIIRLEPFIKHCALAP 462
Rh.s._MdoH   183 GSAYPFAVILDADSLMEGETLVDMVRRMEAEPRLGLLQTLPVVTKARARFGRSMQFSAALHAPVFARGLAMMQGRTGPFWGHNAIVRVQAFAESCGLPE 281
N.e._MdoH    336 GSQYSYMVVLDADSVMSGECLTSLVRLMEANPNAGIIQSWPRASGMDTFYARCQQFATRVYGPLFTAGLHFWQLGESHYWGHNAIIRVQPFIEHCTLAL 434

M.x._MdoH    290 LPGQQPFGGHILSHDFVEAALMRRAGYTVWLVTELGGSFEQPPPSLLDYAQRDRRWCQGNLQHLSLVAAGGLHPISRGHFLMGVMSYAASPLWLLFLMS 388
E.c._OpgH    435 LPGEGSFAGSILSHDFVEAALMRRAGWGVWIAYDLPGSYEELPPNLLDELKRDRRWCHGNLMNFRLFLVKGMHPVHRAVFLTGVMSYLSAPLWFMFLAL 533
P.s._MdoH    446 LPGKGAFAGAILSHDFVEAALMRRAGWGVWIAYDLPGSYEELPPNLLDELKRDRRWCHGNLMNFRLFLVKGMHPVHRAVFLTGVMSYLSAPLWFFFLVL 544
D.c._MdoH    444 LPGEGSFAGSILSHDFVEAALMRRAGWGVWIAYDLPGSYEELPPNLLDELKRDRRWCHGNLMNFRLFLVKGMHPVHRAVFLTGVMSYLSAPLWFMFLAL 542
X.c._MdoH    321 LRGRKPFGGHVLSHDFVEAALMRRGGWAMHMVPYLQGSYEEGPPTLTDLLIRDRRWCQGNLQHAKVVSAKGLHWISRMHMLIGIGHYFTAPMWGLLMLI 419
Ra.s._MdoH   463 IPGKGSLSGEIMSHDFVEAALMRRAGWAVWIAYDLDGSYEEMPPNLLDELGRDRRWCHGNLMNFRLFGSPGFHPVHRAVFVTGVMAYLSAPLWFLFLLL 561
Rh.s._MdoH   282 LSGPPPFGGHVMSHDYVEAALLARAGWIVRFDDDIRGSYEEGPENLVDHAKRDRRWCQGNLQHGRILFAPGLCGWNRFVFLQGIMAYIAPLFWLGFIMA 380
N.e._MdoH    435 LPGEGTFAGSILSHDFVEAALMRRAGWGVWIAYDLPGSYEELPPNLLDELKRDRRWCHGNLMNFRLFLVKGLHPVHRAVFLTGVMSYLSAPLWFMFLAL 533

M.x._MdoH    389 GLIAGLYDGWLSFSNPHLL-EDLGPEA--LAFDTTGAVRLFSVSMAMLFAPKCFGLLLALASSQDSALMGGRVRLMLSVVLESVLATLMAPVMMLFQSH 484
E.c._OpgH    534 STALQVVHAL---TEPQYFLQPRQLFPVWPQWRPELAIALFASTMVLLFLPKLLSILLIWC--KGTKEYGGFWRVTLSLLLEVLFSVLLAPVRMLFHTV 627
P.s._MdoH    545 STALLAVNTL---MEPTYFLEPRQLYPLWPQWHPEKAVALFSTTIVLLFLPKLLSVILIWA--KGAKGFGGKFKVTVSMLLEMLFSVLLAPVRMLFHTR 638
D.c._MdoH    543 STALQVVHTL---MEPQYFLQPRQLYPVWPQWRPELAIALFSTTLVLLFLPKLLSVILVCA--KGAKSYGGTARLCLSLLIEMLFSVLLAPVRMLFHTV 636
X.c._MdoH    420 GIGIPLAGGGIDLAGDLPF----SPARYWHGSSQGNAIWIFICTMFVLLAPKLLGYIALLLNPRELRACGGAFRAAVSILLETVLAALMAPVVMYLQSR 514
Ra.s._MdoH   562 STALLAKHTL---IAPEYFTQPRQLFPIWPEWHPEKAAALFSATATVLFLPKILSVLVLWA--KGPRRFGGALHLALSMVIEAVFSVLAAPVRMLFHTR 655
Rh.s._MdoH   381 SIAAPFFAPPL-----DYFPVPYWPFPVFPSDETWKAIGLAVGIFGLLLLPKLMIAIEAIVT-GRAAGFGGAGRVLISTLAELVFSSIIAPILMAFQTR 473
N.e._MdoH    534 STALQVVHAF---TEPHYFLQPHQLFPVWPQWQPELAIALLASTMVLLFLPKLLSILLIWC--KGTKEYGGFIRVTISLLLEVILSVLLAPVRMLFHTV 627

M.x._MdoH    485 FVFGTLLGYKVTWSSQQRDDADLPWAEATRRHAVHTTVGVVLAAVAFFLSPGLLLWLSPVVAGLLLSIPLSVFTSRASLGLWLERRGLLVIPEETEPPR 583
E.c._OpgH    628 FVVSAFLGWEVVWNSPQRDDDSTSWGEAFKRHGSQLLLGLVWAVGMAWLDLRFLFWLAPIVFSLILSPFVSVISSRATVGLRTKRWKLFLIPEEYSPPQ 726
P.s._MdoH    639 FVLAAFLGWAATWNSPQRDDDSTPWIEAVKRHGPQTLLGACWALLVFWLNPSFLWWLAPIVVSLMLSIPVSVISSRTNLGVKARDEKFFLIPEEFEPPQ 737
D.c._MdoH    637 FVVSAFLGWSVQWKSPQRDDDATPWSEAFARHGSQLLLGLVWAGGMAWLDLRFLWWLGPIVFSLILSPLVSVLSSRAGLGLACKRGKLLLIPEEYEPPR 735
X.c._MdoH    515 GVFEVLAGKDSGWDAQVRDDGKLSWPALLRSYGGLTVFGLFMGAVAYAVSPALAAWMGPVIVGMALSIPVVALTSLRRSGMALRRAGIFCIPEELDPPK 613
Ra.s._MdoH   656 FVTAAFLGWKVHWKSPPRDDAQTHWGDAVRRHGLHTLLGLGWAALVYWLNPSFLWWLSPVVGALIVSILLSVFSSRVTLGRGMRRWRLFMIPEEVRPPR 754
Rh.s._MdoH   474 SVLQVLLGRDGGWPTNNRGDGSLSVAQAWSASHWIVTWGLIGIGATYYFAPGLVPWLLPVALPMIFSPLVIAVTSKRS------RSALFTMPLEVAPTP 566
N.e._MdoH    628 FVVSAFLGWKVAWKSPQRDDDSTTWREAFMRHGSQLLLGLVWATGMAWLDLRFLFWLAPIVFSLILSPFVSVFSSRASVGLRAKRWKLLLIPEEYSPPK 726

M.x._MdoH    584 VLERAQEVAEEHALEPVKDAVDHVISDAKAHALHLALLESQPSPAAVPMALASARRK--LLGDNVEPLSPPEKTAVLMDAPTLTEARERRLTLAVS--- 677
E.c._OpgH    727 VLVDTDRFLEMNRQRSLDDGFMHAVFNPSFNALATAMATARHRA-SKVLEIARDRHVEQALNETPEKLNRDRRLVLLSDPVTMARLHFRVWNSPER--Y 822
P.s._MdoH    738 ELISTDQYTYENRWHALKQGFIRAVVDPRQNALACALATSRHRQ-AQPIEVVRMERVDQALKVGPAKLGNQERLMLLSDPVALGRLHERVWSEGHEEWL 835
D.c._MdoH    736 ELVATDEYFRRNRERKLENGFMHAIFDPSINALVSAMATARHRL-SKPIENMREQRVNEALSRKPQDVDGNLRLALISDPVTLARLHHRVWSQPEN--Y 831
X.c._MdoH    614 VLVRASELRRAAALEPSLI-------------------------------------------------------------------------------- 632
Ra.s._MdoH   755 ELRAMRKHL---RQAPPTPDFRLAVVDPVTNALMCAIGTARFPH-DPKLLEVRDATVLQALAAGPDKLTSKQKLVLLSDPLALSALHLAVWSSDVH--R 847
Rh.s._MdoH   567 VLLAHDAILADWERSPAPEAVPALAVSHA---------------------------------------------------------------------- 595
N.e._MdoH    727 VLVDTDNYLMMNRNRTLNDGFMHAVFHPSFNALTTATATARHRK-SKVLEIARDHHIEQALNEPPDKLNRDCRLTLLSDPVIMSRLHYCVWAMPEK--Y 822

M.x._MdoH        ---------------------------------------------
E.c._OpgH    823 SSWVSYYEGIKLNPLALRKPDAASQ--------------------     847
P.s._MdoH    836 AAWRASIEADPHAPLLPLQPVGKAS------EPVPV---------     865
D.c._MdoH    832 GNWRSDYQMLAAPMIKSE---------------------------     849
X.c._MdoH        ---------------------------------------------
Ra.s._MdoH   848 AAWLPANALA------SKAAA------------------------     862
Rh.s._MdoH       ---------------------------------------------
N.e._MdoH    823 ASWVNHYQQLTLNPSALKQCEPNIEEEADYAEPTAQIDMALPGTP     867
```

**Figure S3:** 1D Proton and 2D COSY NMR spectra of MXP-A.

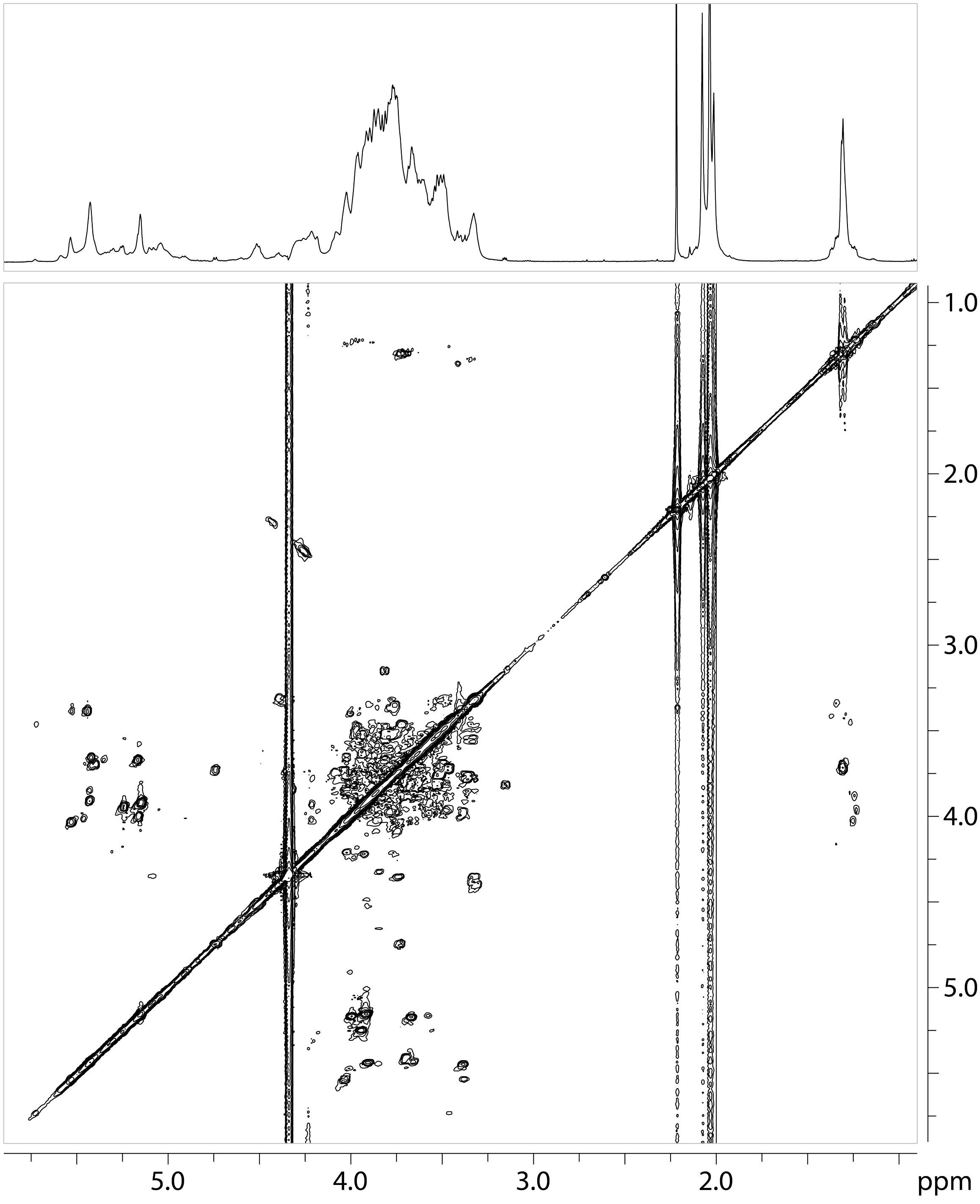

**Figure S4:** Partial 2D NOESY NMR spectrum (*A*) and overlaid HSQC (*B*, black trace), and HMBC (B, magenta trace) NMR spectra of MXP-AS. The letters in the labels refer to individual residues as shown in Figure 3A and Table S2. The numbers refer to the positions of each ring beginning with the anomeric position as Number 1. The red and blue labels indicate the inter-residue HMBC and NOE correlations, respectively. These correlations define the sequence of the polysaccharide. The spin system labeled with the asterisk stems from a residual GlcNAc residue periodate-oxidized on C3 and C4. This fragment was not completely removed by the hydrolysis.

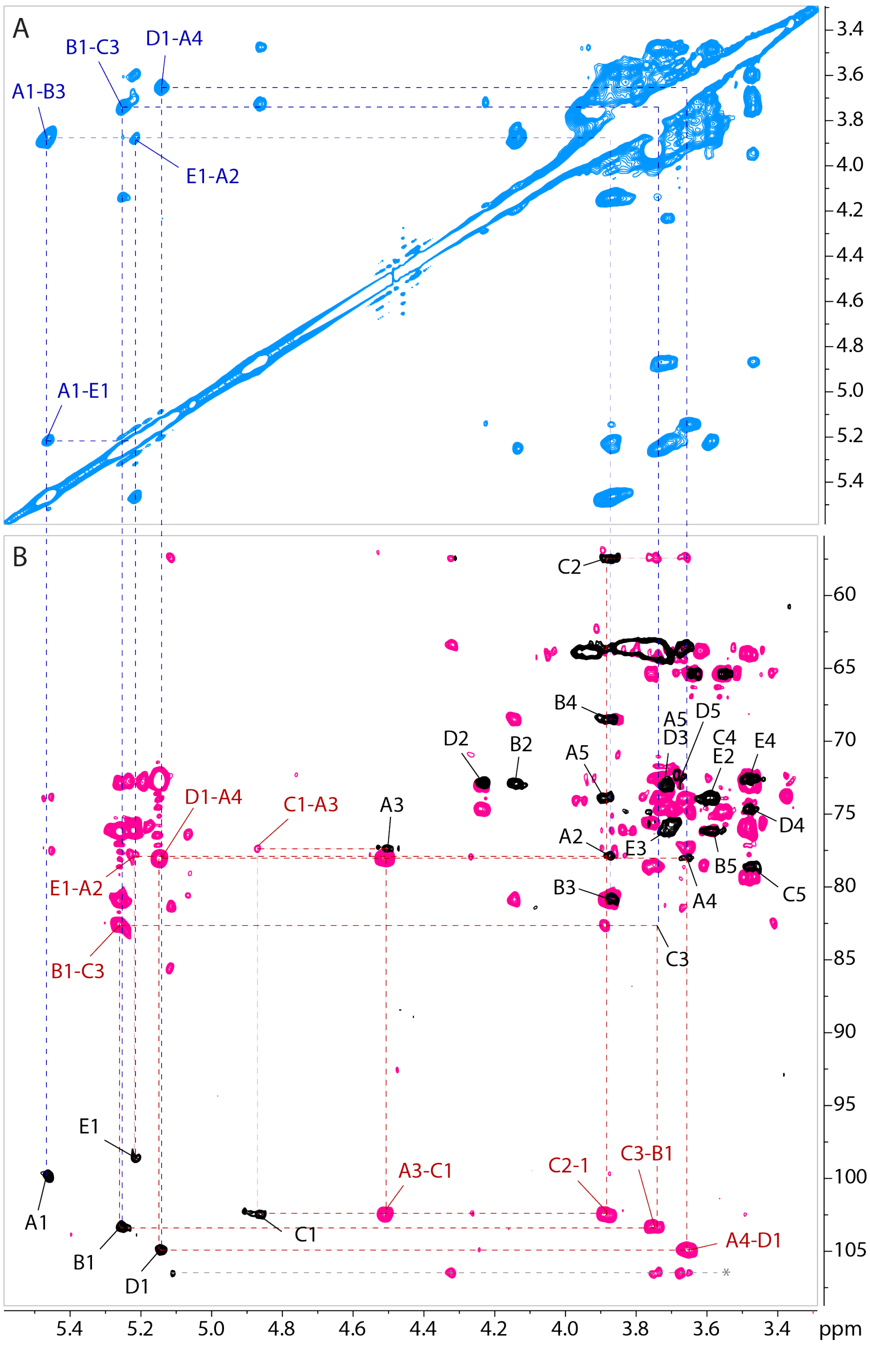

**Figure S5:** Partial 2D NOESY NMR spectrum (*A*) and overlaid HSQC (*B*, black trace), and HMBC (B, magenta trace) NMR spectra of MXP-A. The letters in the labels refer to individual residues as shown in Figure 3C and Table S3. The numbers refer to the positions of each ring beginning with the anomeric position as Number 1. The red and blue labels indicate the inter-residue HMBC and NOE correlations, respectively. These correlations define the sequence of the polysaccharide.

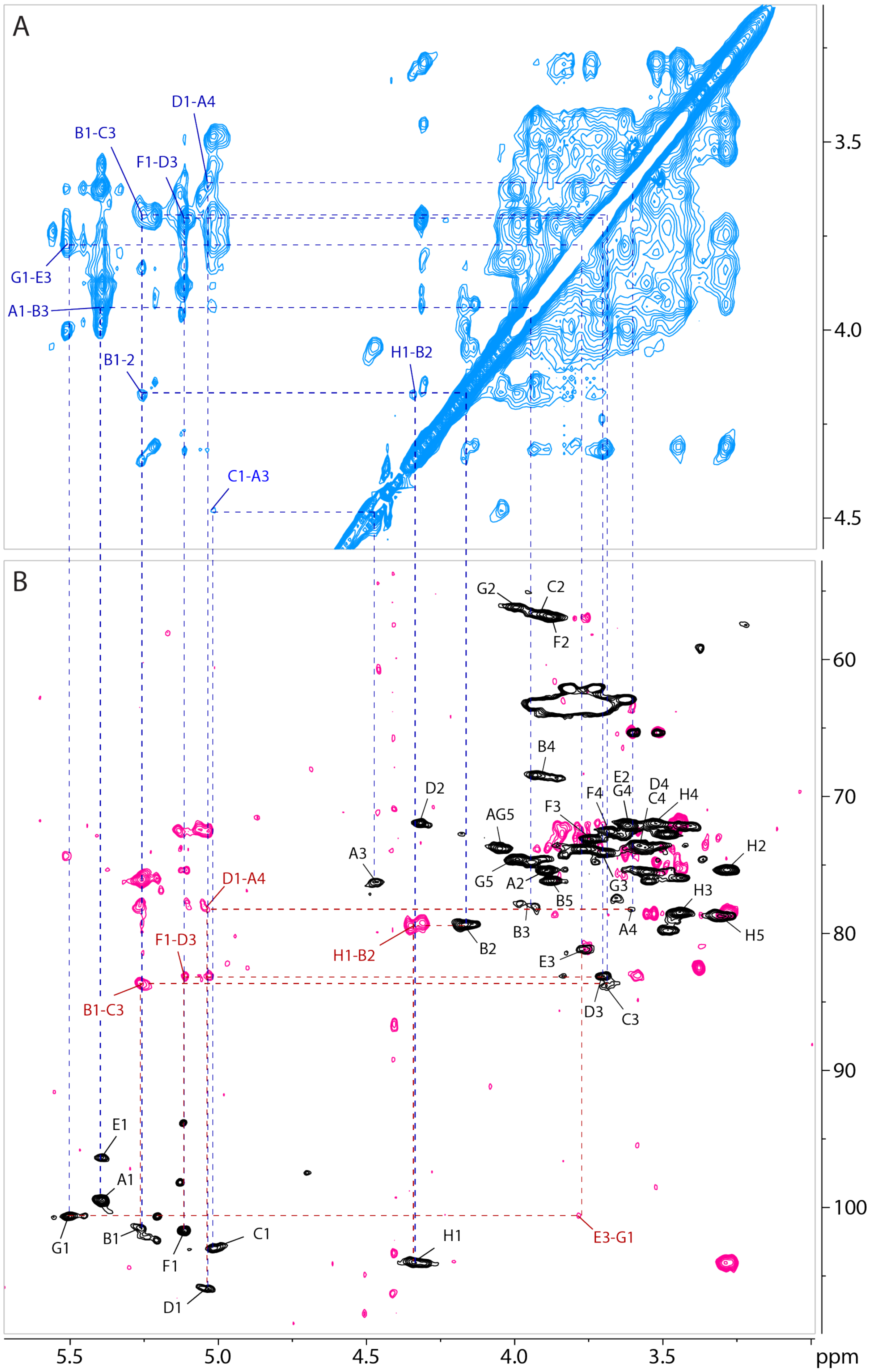
